# Quantifying cancer- and drug-induced changes in Shannon information capacity of RTK signaling

**DOI:** 10.1101/2025.04.30.651439

**Authors:** Paweł Nałęcz-Jawecki, Lee Roth, Frederic Grabowski, Sunnie Li, Marek Kochańczyk, Lukasz J. Bugaj, Tomasz Lipniacki

## Abstract

Signaling pathways transmit and process information, enabling cells to respond accurately to external cues. Disease states like cancer can corrupt signal transmission, though the magnitude to which they reduce information capacity has not been quantified. Here we apply pseudo-random pulsatile optogenetic stimulation, live-cell imaging, and information theory to compare the information capacity of receptor tyrosine kinase (RTK) signaling pathways in EML4-ALK-driven lung cancer cells (STE-1) and non-transformed lung epithelial cells (BEAS-2B). The information rate through the RTK/ERK pathway in STE-1 cells was below 0.5 bit/hour but increased to 3 bit/hour after oncogene inhibition. Information was transmitted by only 50–70% of cells, whose channel capacity (maximum information rate) was estimated through *in silico* protocol optimization. Although oncogene inhibition increased the capacity of the RTK/ERK pathway in STE-1 cells (6 bit/hour), capacity remained lower than in BEAS-2B (11 bit/hour). The capacity of the parallel RTK/calcineurin pathway in BEAS-2B exceeded 15 bit/hour. This study highlights information capacity as a sensitive metric for identifying disease-associated dysfunction and evaluating effects of targeted interventions.

## Introduction

Biochemical signaling pathways are responsible for the regulation of intracellular processes in response to extracellular cues, infections, or cell damage. Ligand stimulation of the RAS/RAF/MEK/ERK pathway through receptor tyrosine kinases (RTKs) (Fig 1A), referred to here as the RTK/ERK pathway, is responsible for cell proliferation, survival, and differentiation (Zhang & Liu, 2002). The RTK/ERK pathway is often stimulated in transient pulses in processes that require high spatiotemporal resolution, such as collective cell migration and embryogenesis (Aoki *et al*, 2017; Hino *et al*, 2020; Corson *et al*, 2003). Mutations that cause constitutive activation of RTK/ERK or PI3K/AKT/mTOR pathways induce uncontrolled cell proliferation and growth (Nishida & Gotoh, 1993; Burotto *et al*, 2014; Song *et al*, 2023; Katso *et al*, 2001), while silencing of other pathways (such as p53 or ATM/ATR) impairs cell cycle arrest and apoptosis in damaged cells (Marei *et al*, 2021; Bartkova *et al*, 2005; Mehrdad & Parvin, 2018), ultimately leading to cancer development. RTKs also trigger calcium signaling (Lee *et al*, 2009), which is crucial for cell cycle control (Berridge, 1995), and changes in calcium activation patterns have been reported in cancers harboring FGFR mutations (Nguyen *et al*, 1997; Katso *et al*, 2001).

**Figure 1.**
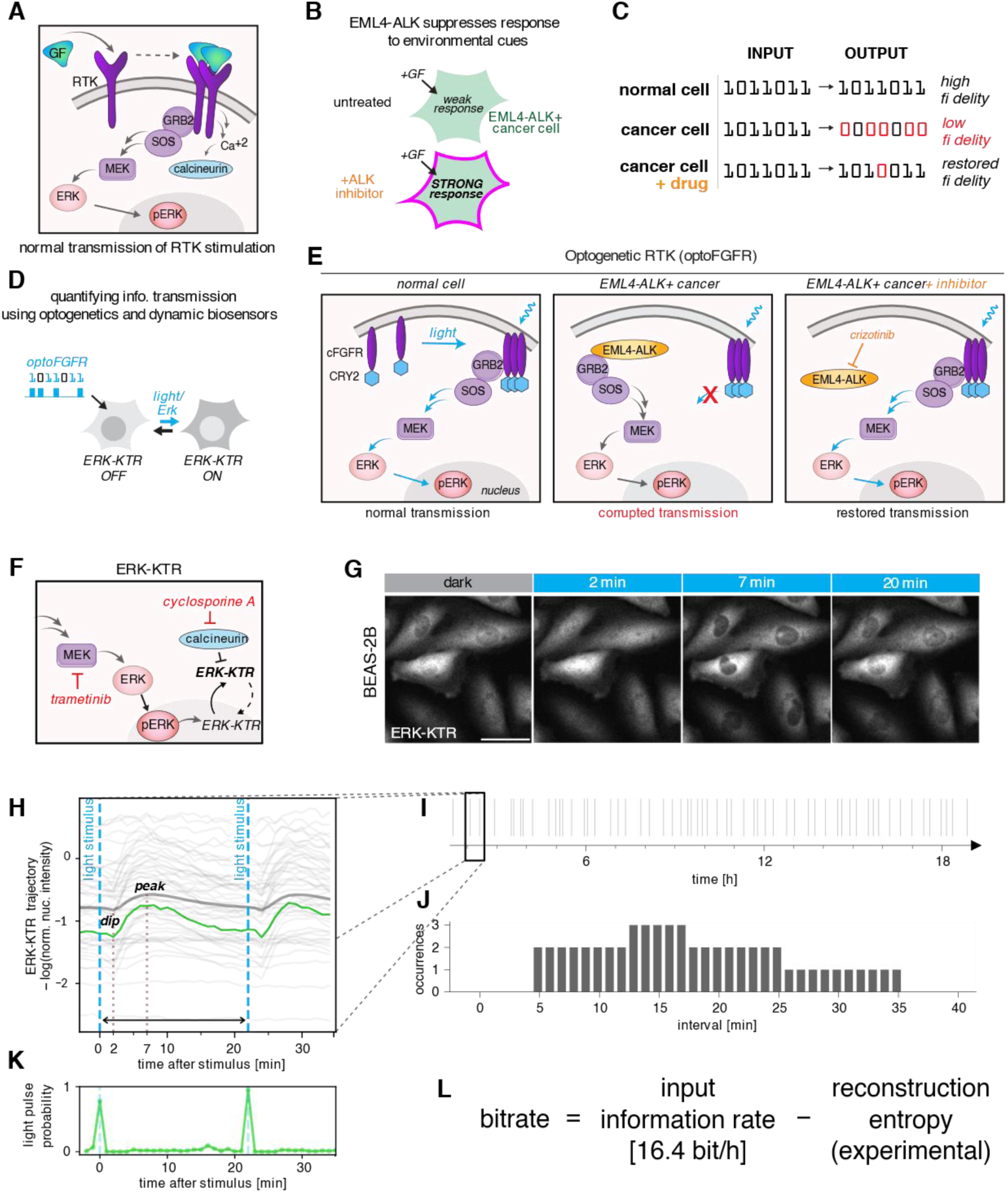
Experimental setup. A After binding to a growth factor (GF), receptor tyrosine kinases (RTKs) multimerize and signal to ERK via RAS, RAF, and MEK (the RTK/ERK pathway). Simultaneously, RTKs trigger the calcium-calcineurin pathway (RTK/calcineurin pathway). B The EML4-ALK oncogene suppresses a cell’s ability to respond to environmental cues like growth factors. ALK inhibitor (ALKi) restores responsiveness. C In normal cells, activity pulses are accurately transmitted by the RTK/ERK pathway from receptors to the nucleus. EML4-ALK-positive cancer cells transmit the environmental information with low fidelity, which can (to some extent) be restored by ALKi. D Pulsatile optogenetic FGFR (optoFGFR) stimulation and fluorescent biosensors of signaling allow quantification of information transmission through the RTK/ERK signaling axis. The ERK-KTR biosensor translocates between the nucleus and cytoplasm upon ERK activation/inactivation. E OptoFGFR, in response to blue light, recapitulates endogenous RTK activation. Cell responsiveness to RTK signals may be suppressed by EML4-ALK, which associates with and sequesters GRB2-SOS. Signal transmission can be restored by ALKi, which liberates GRB2- SOS from EML4-ALK. F ERK-KTR migrates to the cytoplasm upon phosphorylation by ERK, and in the reverse direction after dephosphorylation by calcineurin. Inhibitors of calcineurin (cyclosporine A) or MEK (trametinib) were used to study the two pathways in isolation. G ERK-KTR cytoplasmic–nuclear shuttling after stimulation with a light pulse in BEAS-2B cells. Fluorescence was recorded with a one-minute resolution. H Light stimulus leads to ERK-KTR phosphorylation, resulting in cytoplasmic translocation (peak is observed about 7 min after the light pulse), preceded by a calcineurin-mediated ‘dip’ observed 2 min after the light pulse in BEAS-2B cells. Thin gray lines show single-cell preprocessed ERK-KTR trajectories, thick gray line shows their average. The green line shows a selected single-cell trajectory for which the (posterior) probability of a light pulse is given in panel K. See Methods for details of ERK-KTR trajectories preprocessing. I Cells were stimulated with a pseudo-random series of light pulses. Information is encoded in intervals between subsequent pulses. J The distribution of intervals between stimulation pulses was chosen based on preliminary experiments to maximize the bitrate. K Single-cell ERK-KTR trajectories were used for probabilistic reconstruction of the input signal, i.e., to predict the probability of a light pulse occurrence at each time point throughout the experiment, at a one-minute temporal resolution. L The numerically estimated entropy of the probabilistic reconstruction was used to compute the rate of information transmission through the pathway.

Beyond elevating basal signaling, cancer mutations can also disrupt important features of signal transmission including its amplitude or kinetics, which may reduce the cells’ ability to respond to extracellular signals accurately (Bugaj *et al*, 2018; Gonzalez-Martinez *et al*, 2024; Gerosa *et al*, 2020). One example is the EML4-ALK fusion oncoprotein (Soda *et al*, 2007; Lei *et al*, 2022), which drives constitutive activation of ERK signaling but simultaneously suppresses signaling through RTKs, rendering the cell insensitive to external growth factors. These disruptions can be reversed by targeted drugs, which restore responsiveness to growth factors and permit pulsatile ERK activity (Fig 1B) (Gonzalez-Martinez *et al*, 2024). Although examples of oncogene-induced changes in signal transmission continue to emerge, it is not clear to what extent cancer alterations corrupt a cell’s capacity to perceive its environment because changes in information transmission through dynamic signaling networks have not been quantified.

Shannon information theory (Shannon, 1948) provides a framework for quantitative assessment of the rate at which information can be transferred through a communication channel, such as a signaling pathway. While *mutual information* measures the information that can be inferred about the input signal given the observed response, *information transmission rate* (or *bitrate*) reflects the amount of information transferred through the channel per unit time. The maximum rate at which information can be transmitted through a channel using the best encoding is called *channel capacity*. Mutual information has been used to show that the response amplitude in several important signaling pathways, including the RTK/ERK pathway, transmits only about 1 bit of information about the strength of a stimulus, indicating that these pathways respond in an all-or-nothing manner (Cheong *et al*, 2011; Uda *et al*, 2013). Slightly more information can be inferred from the temporal pattern of the response (Selimkhanov *et al*, 2014). Analysis of information transmitted by single cells has revealed that variability in cellular responses is primarily driven by phenotypic diversity rather than the intrinsic stochasticity of signaling pathways (Tkačik *et al*, 2008; Tudelska *et al*, 2017; Kramer *et al*, 2022; Topolewski *et al*, 2022; Kramar *et al*, 2024). Information transmission rate is an adequate measure in systems where repeated use of a channel rather than a one-time decision is important, because it accounts for both the fidelity of signal transmission and its temporal resolution. In such systems, information may be encoded in the intervals between signaling pulses. Recently, we showed that such encoding allows transmission of at least 6–8 bit/h through the RTK/ERK pathway (Nałęcz-Jawecki *et al*, 2023), exceeding earlier theoretical estimates (Grabowski *et al*, 2019).

In this study, we apply Shannon information theory together with optogenetics and live cell imaging to quantify changes to signal transmission resulting from the EML4-ALK fusion oncoprotein in an EML4-ALK+ cell line, STE-1. We show that in this cell line, information transmission through the RTK/ERK pathway is almost entirely blocked. Treatment with the ALK inhibitor (ALKi) crizotinib restored information transmission, although to a rate still twice lower than that observed in non-cancerous BEAS-2B cells (Fig 1C). In these cells, RTK- triggered calcium/calcineurin signaling transmitted even more information than ERK, although activation of this pathway was not observed in STE-1. Our work demonstrates how information-theory can provide a quantitative, functional metric to understand the oncogenic state and to assess potential therapies.

## Results

### Pulsatile optoFGFR stimulation enables estimation of information flow in the RTK/ERK pathway

To monitor and induce the activity of the RTK/ERK pathway, we engineered cells to express a fluorescent reporter of ERK activity (ERK-KTR) and an optogenetic FGF receptor (optoFGFR), which allows us to stimulate the pathway at precisely selected time points using light (Fig 1D). We investigated an EML4-ALK-expressing human non-small-cell lung carcinoma cell line, STE-1, and a non-cancerous cell line, BEAS-2B, as a reference. In normal cells, such as BEAS-2B, optoFGFR activation triggers RTK/ERK pathway signaling, culminating in ERK phosphorylation/activation and subsequent phosphorylation of its downstream targets. In STE-1 cells, EML4-ALK hijacks GRB2-SOS, leading to constitutive ERK activation and also to active suppression of transmembrane RTKs, including optoFGFR (Gonzalez-Martinez *et al*, 2024). ALK inhibition relieves this suppression and restores the cells’ ability to transmit RTK signals (Fig 1E). ERK activity can be monitored by ERK-KTR, which, when phosphorylated by active ERK, translocates to the cytoplasm (Fig 1F,G). In some cell types, including BEAS-2B, FGFR (and optoFGFR) also triggers calcium signaling (Kim *et al*, 2014), which leads to the activation of the phosphatase calcineurin (Fig 1A). Because ERK-KTR was engineered from a natural substrate for calcineurin (ELK-1), calcineurin can dephosphorylate and inactivate ERK-KTR, driving its nuclear localization (Dessauges *et al*, 2022). To disambiguate signaling via the RTK/ERK and RTK/calcineurin pathways, we performed experiments in the presence or absence of inhibitors of MEK (trametinib, MEKi) and/or calcineurin (cyclosporine A, CALCi; Fig 1F). In a typical experiment, 100 ms light pulses were administered at prespecified intervals (see below), and ERK-KTR fluorescence (Fig 1G) was recorded every minute, resulting in single-cell trajectories of ERK-KTR nuclear intensity. In BEAS-2B cells, the preprocessed ERK-KTR trajectories (see Methods for preprocessing details) typically featured a small ‘dip’ 2 min after stimulation, resulting from the calcineurin-mediated ERK-KTR dephosphorylation. This was followed by a cytoplasmic ERK-KTR translocation peak 7 min after the light pulse, mediated by the RTK/ERK pathway (Fig 1H).

To measure the rate of information transmission, we stimulated the STE-1 and BEAS-2B cells with a pseudo-random series of light pulses (Fig 1I). The intervals between subsequent light pulses followed a fixed distribution (Fig 1J), carefully chosen based on preliminary experiments to ensure both near-optimal bitrate and sampling of a broad range of intervals. Single-cell ERK-KTR trajectories enabled a probabilistic, machine learning-based reconstruction of the input signal (Fig 1K). We used this reconstruction to compute the entropy *H*(*X* | *Y*) of the input conditioned on the observed response (i.e., information lost due to the uncertainty of signal reconstruction). The amount of transmitted information is then given by *I*(*X*; *Y*) = *H*(*X*) − *H*(*X* | *Y*), where *H*(*X*) is the input entropy (sent information). Therefore, we computed the transmitted information rate as the input information rate minus the reconstruction entropy rate (Fig 1L). Preliminary experiments on BEAS-2B and STE-1 cells, along with an earlier study on MCF-10A cells (Nałęcz-Jawecki *et al*, 2023), indicated that the effective refractory time of the RTK/ERK pathway is approximately 8–10 min. The chosen interval distribution includes intervals from 5 to 35 min and has an input entropy rate of 16.4 bit/h. In principle, the input information rate could be higher for more frequent pulses, reaching 60 bit/h for pulses occurring every minute with a probability of ½. However, if light pulses were sent at intervals shorter than the refractory time, the cell responsiveness would deteriorate, increasing the reconstruction entropy.

### EML4-ALK blocks information transmission, which can be restored by ALK inhibition

To estimate the reconstruction entropy (and thus the amount of information transmitted by the sequence of light pulses), we trained a multi-layer perceptron (MLP with 3 layers) using all single-cell tracks longer than 3 hours from both cell lines, all experimental conditions and all experimental replicates (see Methods for details). To predict the probability of a light pulse at a given time (Fig 1K), the MLP used a 7-min fragment of the ERK-KTR trajectory, the time elapsed since the previous light pulse, a measure of the track variability, and information about the cell line and an optional inhibitor. The last two inputs allowed a single MLP to generalize across experiments in which cells follow qualitatively different trajectories. In training, we directly minimized the cross-entropy, constituting an upper bound of the conditional entropy *H*(*X* | *Y*). By subtracting the result from the input entropy *H*(*X*), we obtained a lower bound on the transmitted information *I*(*X*; *Y*). We noticed that cells with low (or very high) expression of optoFGFR have markedly lower bitrate, and therefore excluded them from further analysis (see Methods and Fig S1). We used the remaining cells (without retraining the MLP) to estimate the population-averaged bitrate at which the STE-1 and BEAS-2B cells transmit information when stimulated according to the selected protocol (Fig 1J).

Consistent with our earlier results (Gonzalez-Martinez *et al*, 2024), we found that without inhibitors, STE-1 cells were unresponsive to light stimulation (Fig 2A,B row 1) and thus transmitted nearly no information from the activated receptor to ERK-KTR (Fig 2C row 1). In contrast, when treated with ALKi, a significant fraction of cells evoked an ERK-KTR trajectory peak (Fig 2A,B rows 2–4), which allowed for an average information transmission rate of ∼3 bit/h (Fig 2C, rows 2–4). The effect did not depend on the ALKi concentration in the tested range (0.3–3 μM), which indicates that the concentration of 0.3 μM is sufficient to maximally restore information transmission. The increase in responsiveness was largely due to a drug-induced decrease in basal signaling, which increased the dynamic range of stimulation (Fig S3).

**Figure 2.**
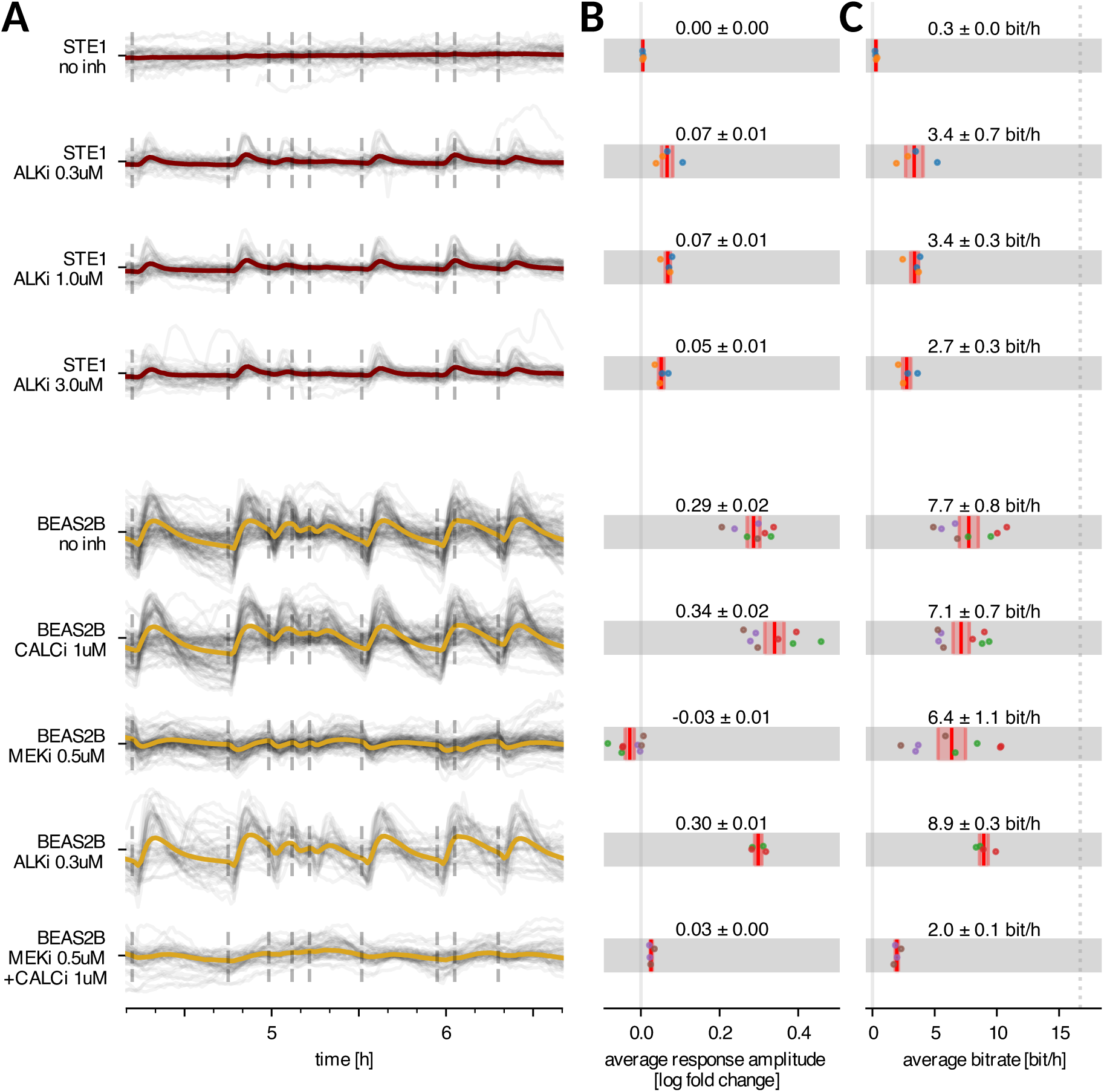
Response amplitude and bitrate in the STE-1 (cancerous) vs. BEAS-2B (non-cancerous) cell lines, with or without inhibitors. A 2.5-hour fragments of ERK-KTR trajectories, obtained in experiments with STE-1 and BEAS-2B cells with or without inhibitors, as indicated. Gray lines show single-cell trajectories, while colored lines represent the population average. Dashed vertical lines indicate light pulses. B Population-averaged response amplitude measured 7 min after light pulse (average is taken over all cells and light pulses; see Fig S2 for response amplitude definition). C Population-averaged bitrate. The dotted line shows the input information rate of the input pulse sequence. In panels B and C, dots correspond to experimental/technical replicates; dots of the same color denote results from the same experimental replicate; bars correspond to the mean and standard error of the mean. Cell lines and stimulation protocols correspond to those in panel A.

We performed the same analysis for BEAS-2B cells, obtaining a 4–6 times stronger average response and a more than twice higher bitrate for the RTK/ERK pathway (7.2 bit/h; Fig 2B,C, row 6), matching results obtained for MCF-10A cells in Figure 4 in (Nałęcz-Jawecki *et al*, 2023). As expected, ALKi minimally affected bitrate in BEAS-2B cells, consistent with its lack of ALK kinase expression. (Fig 2C, row 8 vs. row 5)

### In BEAS-2B cells, the RTK/ERK and RTK/calcineurin pathways transmit information independently

As mentioned before, optoFGFR signals to ERK-KTR not only through ERK but also through calcineurin. The latter pathway is faster and evokes a dip (nuclear ERK-KTR translocation) in the ERK-KTR trajectory (Fig 1H), preceding the peak (cytoplasmic ERK-KTR translocation) induced by ERK. To verify whether the high bitrate observed in BEAS-2B cells results from the MLP classifier using this dip to detect light pulses, we inhibited the RTK/calcineurin pathway with a CALCi. While CALCi eliminated the dip and slightly increased the response amplitude (Fig 2B, row 6), it only marginally reduced the bitrate, indicating that the RTK/ERK pathway alone can transmit approximately 7.2 bit/h (Fig 2C row 6). Similarly, after blocking the RTK/ERK (RTK-RAS-RAF-MEK-ERK) pathway with MEKi, the RTK/calcineurin pathway was capable of transmitting approximately 6.4 bit/h. When operating simultaneously (without any inhibitors), the two pathways transmit 7.8 bit/h. As a control, we confirmed that the joint application of CALCi and MEKi prevents effective information transmission, reducing the average response amplitude tenfold and the bitrate to 2.1 bit/h.

Overall, these results show that in the non-cancerous cell line, both the RTK/ERK and the RTK/calcineurin pathway, separately or jointly, transmit around 7 bit/h when stimulated according to the chosen encoding protocol. Information transmission is fully suppressed in STE-1 cells expressing EML4-ALK, but can be restored to 3 bit/h by ALKi treatment.

### In information-transmitting cells, the bitrate of ALKi-treated STE-1 cells remains half that observed in BEAS-2B

To explore the differences between STE-1 and BEAS-2B cells in more detail, we assessed the heterogeneity of individual cell bitrates within both cell lines. As illustrated in Fig 3A, the bitrate distributions spanned a wide range, from approximately −2.5 bit/h to 16.4 bit/h (the upper bound equals the input entropy rate). Negative bitrates are possible and indicate that a particular cell responded misleadingly (with respect to the majority of cells in the population), as explained in the Bitrate estimation section in Methods. In BEAS-2B cells, the histograms exhibit a distinct bimodal pattern: one group of cells clusters near zero, while the other centers around 12–13 bit/h. In STE-1 cells with ALKi, the two modes overlap, with only a tiny fraction of cells transmitting more than 10 bit/h. This shows that even with ALKi treatment, STE-1 cells rarely transmit information at rates characteristic of transmitting BEAS-2B cells.

**Figure 3.**
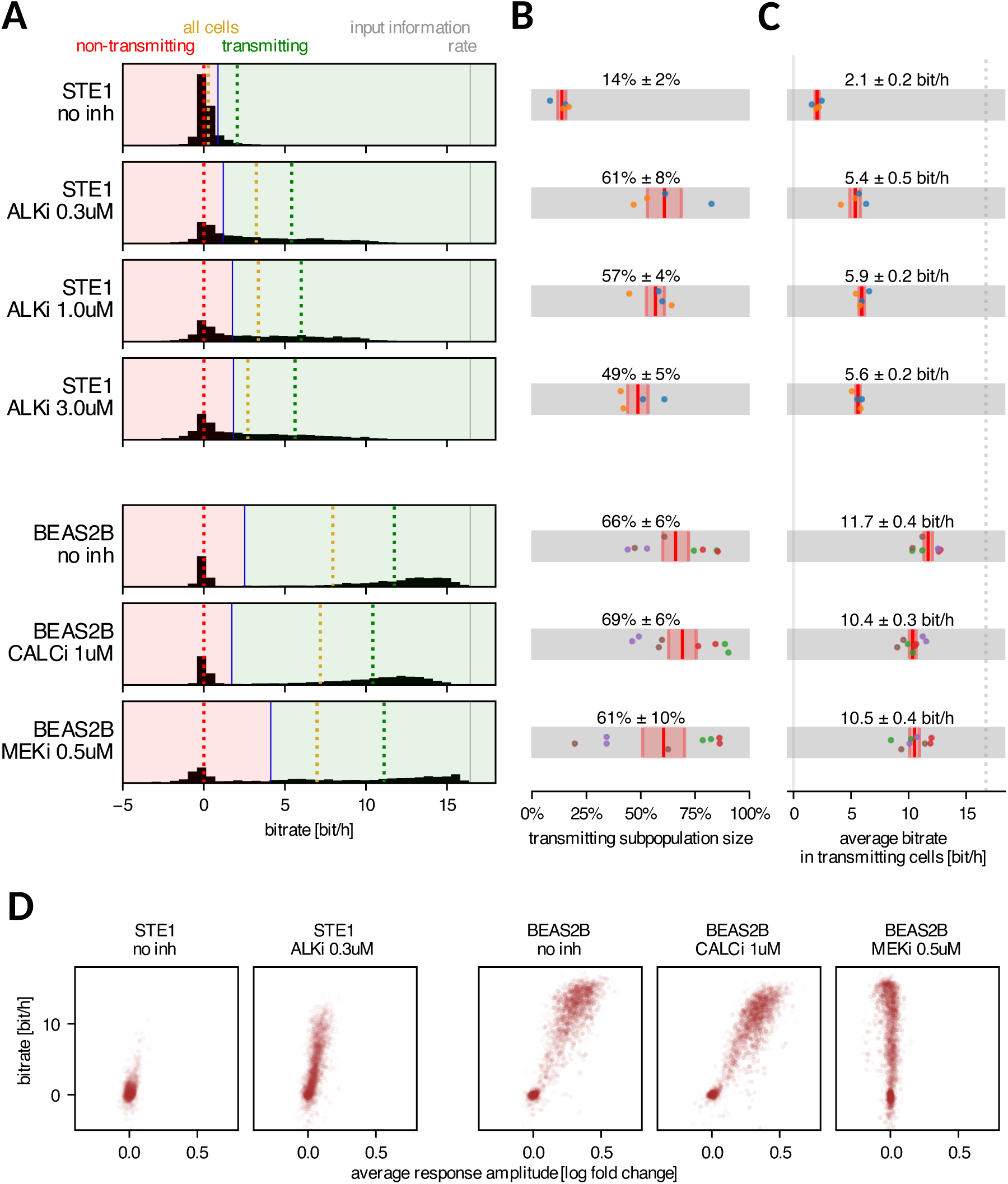
Bitrate of individual cells. A Histograms of bitrates of individual cells; for each condition, all biological and technical replicates are pooled. Cells were divided into transmitting and non-transmitting subpopulations so that the average bitrate in the non-transmitting subpopulation is zero. Dotted lines show bitrate averaged over non-transmitting (red), transmitting (green), and all cells (yellow). B Fractions of transmitting cells computed for each replicate separately; average and standard error of the mean are shown. C Average bitrate in the transmitting cell subpopulation computed for each replicate separately; average and standard error of the mean are shown; dotted line indicates the input information rate. D Average response amplitude and bitrate in individual cells; dot intensity is proportional to the duration each cell was tracked. In panels B and C, dots correspond to experimental replicates; dots of the same color denote results from the same experiment. The threshold shown in panel A (blue line) was computed for all replicates pooled, whereas in panels B and C, the threshold was chosen for each replicate separately.

Formally, we split the cells into ‘non-transmitting’ and ‘transmitting’ subpopulations by setting a threshold on the histogram such that cells to the left of this threshold (‘non-transmitting’) transmit on average 0 bit/h. The average fraction of transmitting cells ranged between 60– 70% for BEAS-2B cells, regardless of applied inhibitors, and between 50–60% for ALKi-treated STE-1 cells (Fig 3B). Untreated STE-1 cells consistently contained approximately 15% transmitting cells.

We noticed that both the average bitrate for the entire cell population (Fig 2B) and the fraction of transmitting cells (Fig 3B) were sensitive to experimental replicate-dependent factors. However, their ratio—the average bitrate within the transmitting subpopulation (Fig 3C)—was much more stable. The transmitting subpopulation of STE-1 cells undergoing ALKi treatment consistently achieved a bitrate of around 5.5 bit/h, while transmitting BEAS-2B cells reached 10–11 bit/h using either the RTK/ERK or RTK/calcineurin channel, and nearly 12 bit/h when neither channel was blocked by inhibitor.

We observed a clear correlation between the response amplitude and the bitrate in individual cells (Fig 3D), indicating that the ‘non-transmitting’ cells are primarily non-responding. This suggests that the higher bitrate observed in BEAS-2B cells, compared to ALKi-treated STE-1 cells, is mainly due to the higher average response amplitude, which allows for better separation of the ERK-KTR peak from background noise.

It is important to emphasize that while the difference in response amplitude between BEAS-2B and treated STE-1 cells is evident, its origin is not obvious. The difference may be associated with cell type or result from richer (2% FBS) serum used for STE-1 to prevent cell death over the 17-hour experimental period.

### *In silico* protocol optimization indicates that RTK/ERK channel capacity is 5–15% higher than the experimentally estimated bitrate

The response amplitude (measured 7 min after the last light pulse) depends on the cell line and generally (except for BEAS-2B cells with MEKi) increases with the time that has passed since the previous light pulse (Fig 4A). We observed the highest response amplitude in BEAS-2B cells without inhibitors or with CALCi. In STE-1 cells without ALKi the response amplitude was close to zero (Fig 2B). With ALKi treatment, STE-1 cells began responding to light pulses following intervals longer than 10 min, but the response amplitude remained consistently smaller than in BEAS-2B cells with CALCi or untreated (Fig 4A). The response amplitude in BEAS-2B cells with MEKi was negative, as the RTK/calcineurin pathway leads to ERK-KTR dephosphorylation and translocation to, rather than from, the nucleus.

**Figure 4.**
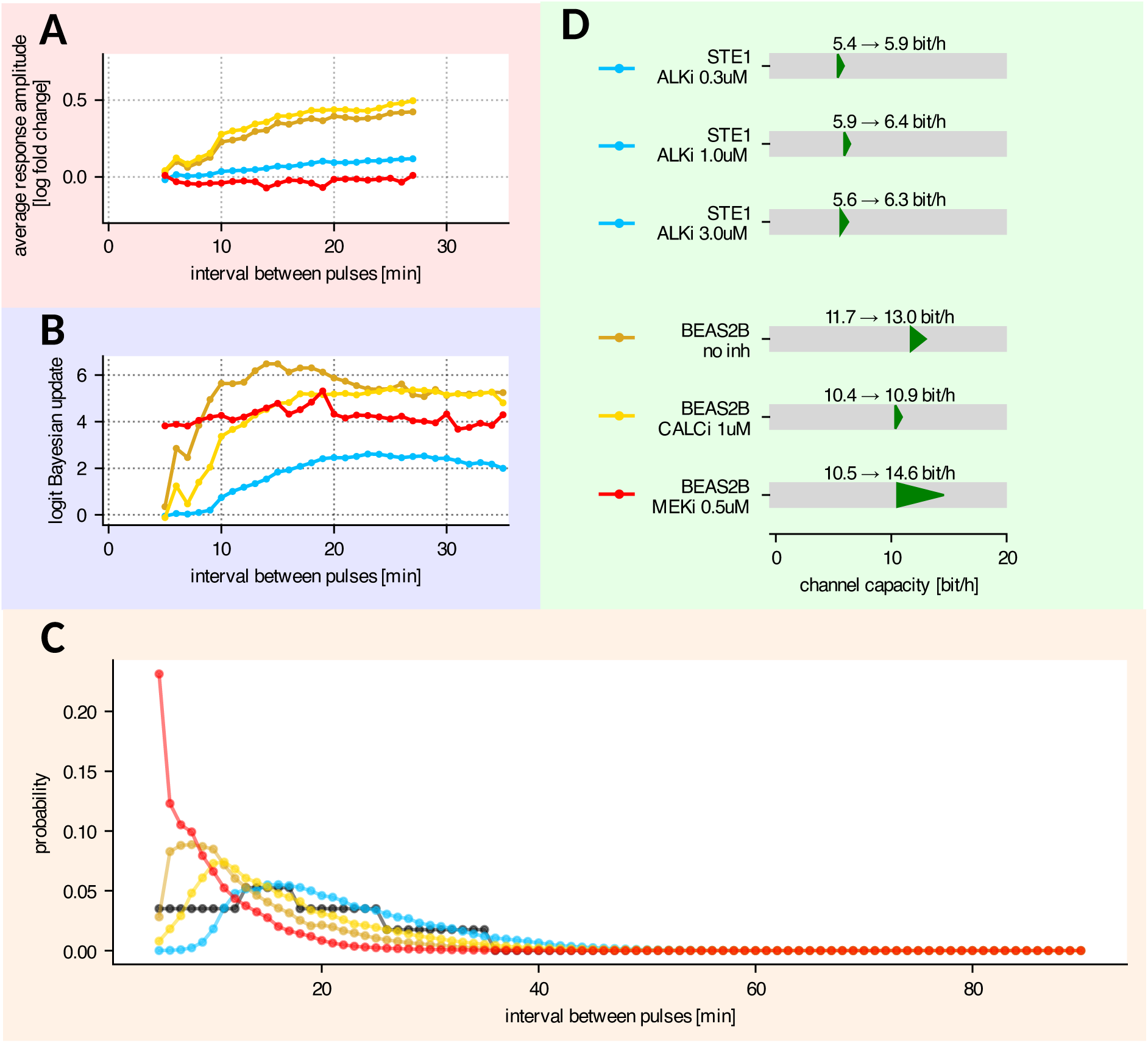
*In silico* protocol optimization and channel capacity estimation. A Average response amplitude 7 min after pulse as a function of interval preceding the light pulse. Colors represent different cell lines and experimental conditions, as indicated in panel D. B MLP classifier’s logit Bayesian update to the prior pulse probability, as a function of interval preceding the light pulse. Colors as indicated in panel D. C Optimized input interval distribution. Colors as indicated in panel D. Black line shows the interval distribution used in the experimental protocol. D Channel capacity (i.e., bitrate achievable with the optimized input interval distribution) in transmitting cells, compared to the experimentally determined bitrates reported in Fig 3C. Data in panels B–D computed based on transmitting cells only. In panels A–D, data from all experimental replicates for a given cell line and condition were pooled and analyzed jointly. In A–C, experiments with three concentrations of ALKi were also considered jointly.

Moreover, the amplitude was only weakly dependent on the time since the previous light pulse, suggesting that the RTK/calcineurin pathway has a very short refractory time. When the response amplitude was measured 2 min after the light pulse (Fig S4), the point at which the calcineurin-induced ERK-KTR dip was the strongest, we observed that both untreated BEAS-2B and BEAS-2B with MEKi responded to intervals as short as 5 min, indicating that the refractory time of the RTK/calcineurin pathway is below 5 min.

Consistent with earlier observations, the certainty of pulse detection by the MLP classifier (Fig 4B) strongly depends on the response amplitude (Fig 4A). The certainty can be measured as the mean logit Bayesian update to the predicted light pulse probability. The logit update is zero when the classifier cannot infer any additional information from the ERK-KTR trajectory, and thus predicts the probability of light pulse equal to the prior probability. In contrast, the logit update approaches the positive or negative infinity when the pulse is, respectively, predicted or excluded with 100% certainty. Detection of light pulses was most reliable in untreated BEAS-2B following intervals longer than 15 min. For shorter intervals, the detection worsened, with light pulses following 5 min intervals being nearly undetectable. When CALCi was applied to BEAS-2B cells, detection of light pulses following intervals shorter than 20 min was worse than in untreated cells, indicating that, in untreated cells, for short intervals the network used both the ERK-mediated peak and the calcineurin-mediated dip to identify light pulses. Light pulse detection based solely on the RTK/calcineurin pathway (in BEAS-2B with MEKi) did not depend on the time since the previous pulse, consistent with the observation that the RTK/calcineurin pathway’s refractory time is likely shorter than 5 min. In STE-1 cells with ALKi treatment, detection of light pulses following intervals of at least 10 min was possible, and the certainty reached saturation at intervals of approximately 20 min duration, similar to BEAS-2B with CALCi. However, even at saturation, the neural network’s detection certainty was markedly lower in STE-1 cells compared to BEAS-2B cells.

Throughout our work, we assume that the length of intervals between subsequent pulses constitutes the ‘alphabet’ for information encoding; particular interval lengths (‘letters’) are independent, and thus the bitrate depends only on the frequency of particular intervals. So far, we estimated the bitrate assuming a fixed distribution of intervals shown in Fig 1J. Although the interval distribution was chosen based on preliminary data to yield a high bitrate for the RTK/ERK channel, it does not have to be optimal for both cell lines and more importantly for the RTK/calcineurin pathway. In contrast to our previous work (Nałęcz-Jawecki *et al*, 2023), the bitrate estimation algorithm employed in this study allowed us to use existing experimental data to make bitrate predictions for protocols that have not been experimentally tested (see Methods). Since the channel capacity is the maximum bitrate that can be achieved by optimizing the encoding protocol, this allowed us to estimate the channel capacity of the investigated pathways *in silico*, without the need to conduct multiple experimental trials. Specifically, for each cell line and inhibitor (all STE-1 experiments with ALKi were pooled), we searched for the optimal input protocol by numerical optimization of the input interval distribution using gradient descent.

We optimized over protocols with interval lengths from 5 min to 90 min. To allow for intervals longer than 35 min (the longest interval in our dataset), we extrapolated cell responses based on the 35 min interval, which was justified by the detection certainty reaching saturation after approximately 20 min (Fig 4B, see Methods for details). In most conditions, cells did not respond to pulses following a 5-min long interval, and thus we assumed that shorter intervals might be safely excluded from the optimized protocols. The only exception was BEAS-2B with MEKi, for which the data presented in Fig 4A,B suggests that pulses following intervals shorter than 5 min may be detectable; consequently, the true channel capacity of the RTK/calcineurin pathway is likely higher than estimated here.

Most of the optimal interval distributions (Fig 4C) follow a qualitatively similar pattern. Intervals below the detectability threshold are not sent; the interval probability rises steeply following the increase in pulse amplitude and detection certainty (Fig 4A,B). Once the detection certainty has reached a plateau, the interval probability starts to decrease, because the usage of long intervals requires more time to relay the same amount of information. The optimal trade-off between the sent information content and time budget per pulse is granted by the exponentially decreasing distribution, which we observe in all optimal protocols. In most conditions, the change in interval distribution due to optimization increased the bitrate by 5–15% (Fig 4D). The only exception was BEAS-2B with MEKi, which experienced nearly 40% gain in the bitrate. In this case, the optimal distribution was significantly shifted towards shorter intervals, which did not affect pulse detection certainty (roughly independent of interval length; Fig 4B), but increased the input information rate.

Overall, we found that ALKi treatment in STE-1 cells, restores the RTK/ERK channel capacity to approximately 6.0–6.5 bit/h. However, this remains below the RTK/ERK channel capacity of about 11 bit/h estimated for BEAS-2B. The RTK/calcineurin channel in BEAS-2B was found to have an even higher capacity of nearly 15 bit/h. A bit surprisingly, this is more than BEAS-2B capacity of transmitting signals from optoFGFR to ERK-KTR when both the RTK/calcineurin and the RTK/ERK pathways are unblocked. This effect, however, can be expected from Fig S4, showing that the absolute value of response amplitude at 2 min after pulse is the highest when MEKi is applied.

## Discussion

The rate at which information is transmitted by signaling channels constrains the complexity of processes that can be regulated within a given time. High bitrates are expected for signaling pathways that regulate cellular responses to rapidly varying extracellular cues. The RTK/ERK pathway, which governs proliferation, differentiation, apoptosis, and motility, is one such pathway, and due to the nature of these processes, its malfunction is frequently associated with cancer.

In this study, using pulsatile optoFGFR stimulation, the presence of the EML4-ALK oncogene in STE-1 cells suppressed RTK/ERK signaling, reducing the pathway’s bitrate nearly to zero. However, treatment with an ALK inhibitor restored information transmission. With the inhibitor, the transmitting subpopulation of cells—comprising approximately 50–60% of all STE-1 cells—achieved an average bitrate of ∼5.5 bit/h. This remained lower than that of the RTK/ERK pathway in non-cancerous BEAS-2B cells, where a transmitting subpopulation of approximately 70% reached an average bitrate of ∼10.4 bit/h. We note that the fraction of transmitting cells and the average bitrate across the entire population exhibited greater variability between replicates than the average bitrate measured within the transmitting fraction. Therefore, we consider the latter a more reliable metric. High correlation between average ERK-KTR response amplitude and bitrate measured in single cells suggests that the ∼5 times higher response strength observed in BEAS-2B cells, compared to STE-1 cells with ALK inhibitor, facilitates pulse detection and is the primary cause of higher bitrate.

In BEAS-2B cells (as in MCF-10A cells, (Nałęcz-Jawecki *et al*, 2023)), two pathways activated by optoFGFR transmit opposing signals to ERK-KTR: the RTK/ERK pathway and the RTK/calcineurin pathway. Calcineurin responsiveness occurs because ERK-KTR contains an ERK docking site from a transcription factor ELK-1 that is also a substrate for calcineurin dephosphorylation (Sugimoto *et al*, 1997; Dessauges *et al*, 2022). The RTK/calcineurin pathway thus induces rapid ERK-KTR dephosphorylation and nuclear translocation, starting 2 min after light pulse stimulation (of note, this rapid nuclear ERK-KTR translocation was not observed in STE-1). In contrast, the RTK/ERK pathway leads to ERK-KTR phosphorylation and cytoplasmic translocation, peaking approximately 7 min after the light pulse. Consequently, in BEAS-2B cells without inhibitors, ERK-KTR trajectories after a light pulse display a small ‘dip’ followed by a more pronounced ‘peak.’ By applying either a calcineurin or a MEK inhibitor, we investigated these pathways in isolation and estimated that each transmits information at a similar rate of 10–11 bit/h (again, restricting estimation to the transmitting subpopulations). These rates are only slightly lower than the bitrate achieved without inhibitors. Our data additionally indicate that, in response to RTK stimulation, ELK-1 might undergo rapid dephosphorylation followed by phosphorylation; the biological meaning of such regulation remains for further study.

The pulsatile stimulation protocol was designed to maximize the bitrate of the RTK/ERK pathway based on preliminary experiments and our previous study on MCF-10A cells (Nałęcz-Jawecki *et al*, 2023). The optimal protocol balances high information input, achieved with short intervals between stimulation pulses, against signal transmission accuracy, which deteriorates when intervals become as short as the refractory time. Since the refractory time varies between cell lines and appears to be much shorter for the RTK/calcineurin pathway, actual information capacities may exceed the measured bitrates. To address this, we developed an approach for optimizing stimulation protocols and estimating channel capacities from a single suboptimal experiment. This method works best when all intervals expected in the optimal protocol are present in the suboptimal protocol. If not, as in our case, responses to the missing intervals must be extrapolated. Our data suggest that this extrapolation is generally justified, with the exception of the experiment using a MEK inhibitor to study the RTK/calcineurin pathway in BEAS-2B cells, where excluding intervals shorter than 5 min may lead to underestimating the channel capacity. The estimated RTK/ERK channel capacities in BEAS-2B cells and STE-1 cells (with ALK inhibitor) are between 5– 15% higher than their experimentally measured bitrates. However, for the RTK/calcineurin pathway, the difference is much greater—approximately 40%. The high RTK/calcineurin channel capacity (∼15 bit/h) results from its short refractory time, allowing the transmission of sequences with short intervals. This estimate should be considered a lower bound, as even higher bitrates might be achieved with stimulation intervals shorter than 5 min. Indeed, calcium signaling plays a role in regulating rapid cellular processes, such as actin remodeling during cell movement or fertilization (Gilkey *et al*, 1978; Whitaker, 2006; DerMardirossian, 2024).

In summary, we developed a machine learning-based method to analyze information flow in signaling channels and its changes as a function of an oncogenic state. We found high information capacities that confirm that these pathways can efficiently transmit signals from receptors to effector proteins and further highlight the impaired transmission in the cancer cells, even in the drug-restored state. Further studies will address whether such high bitrates are fully utilized in physiological responses, whether their suppression plays a role in cancer development, and whether drug-induced relief of suppression plays an important role during therapy, for example in drug resistance.

## Materials and Methods

### Reagents and Tools table

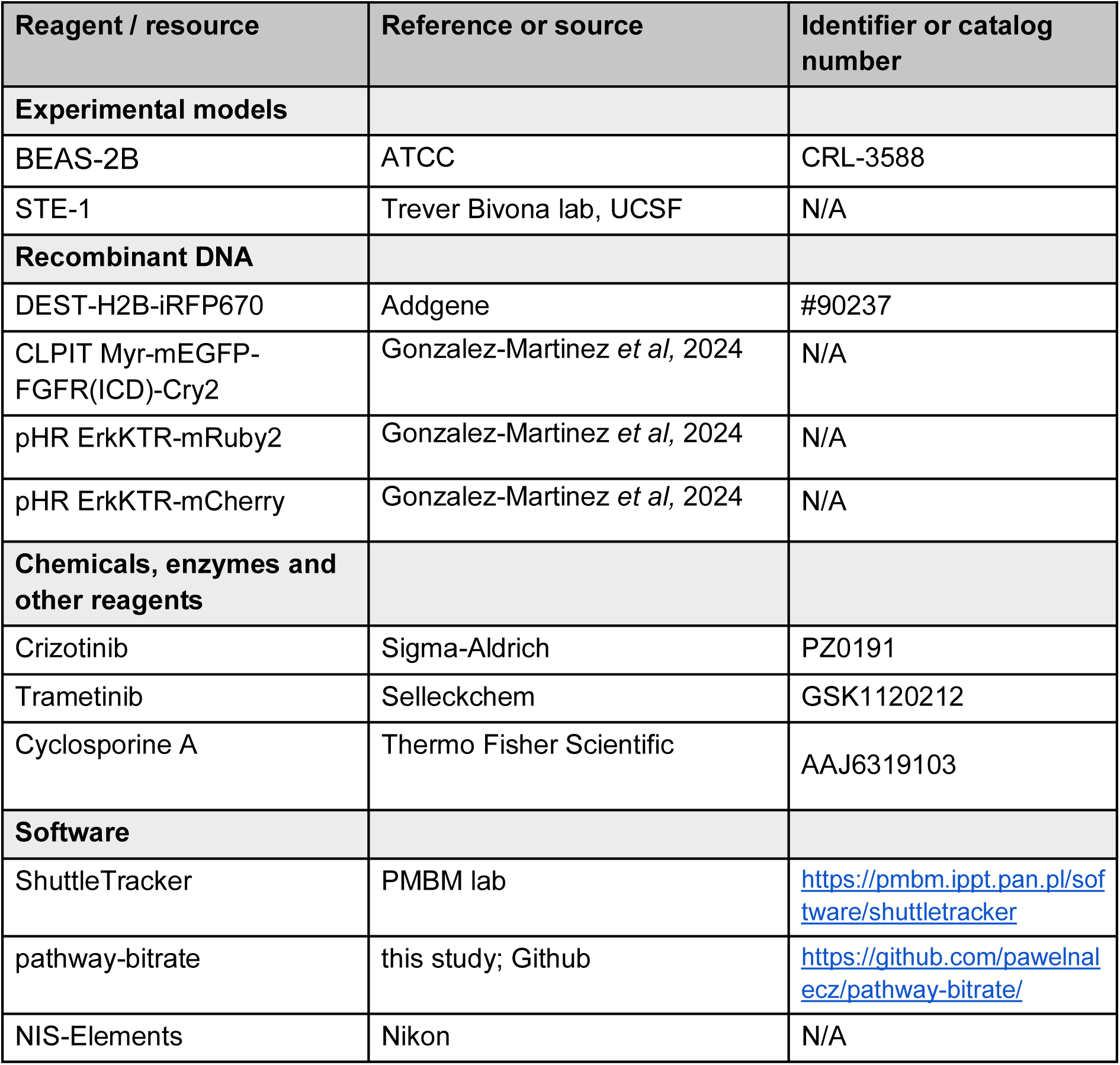

### Methods and Protocols

#### Cell lines and culture

BEAS-2B and STE-1 cells were cultured at 37°C and 5% CO_2_ in RPMI-1640 growth medium supplemented with 10% fetal bovine serum (FBS) and 1% penicillin/streptomycin (P/S). For experiments, cells (STE-1, 1.5×10^4^; BEAS-2B, 5×10^3^) were seeded in 96-well plates (Cellvis) coated with fibronectin (MilliporeSigma, FC01010MG) diluted to 10 µg/mL in PBS. Media were replaced 16 hours before the experiment with RPMI-1640 w/o phenol red, either serum-free or containing 4% FBS (for BEAS-2B or STE-1, respectively). Upon initiation of experiment, equal volume of serum-free media with supplements or DMSO was added to reach the indicated concentrations. Crizotinib (ALKi, Sigma-Aldrich, PZ0191), trametinib (MEKi, Selleckchem, GSK1120212), cyclosporine A (CALCi, Thermo Fisher Scientific, AAJ6319103).

For live cell tracking we visualized cell nuclei using pLentiPGK DEST-H2B-iRFP670 (Addgene:#90237). CLPIT Myr-mEGFP-FGFR(ICD)-Cry2 was cloned by substitution of mCherry with mEGFP in CLPIT Myr-mCherry-FGFR(ICD)-Cry2, infected to cells with pHR ErkKTR-mRuby2 or pHR ErkKTR-mCherry (STE-1 or BEAS-2B, respectively). Expression of these vectors in combination and separately in these cell lines was previously described (Gonzalez-Martinez *et al*, 2024). Infected cells were sorted twice to enrich for high expression of all 3 markers.

Live-cell imaging was performed using a Nikon Ti2-E microscope equipped with a Yokagawa CSU-W1 spinning disk, 405/488/561/640 nm laser lines, an sCMOS camera (Photometrics), and a motorized stage. Cells were maintained at 37°C and 5% CO_2_ using an environmental chamber (Okolabs). Cells were imaged every 60 seconds for 90 minutes without optoFGFR stimulation followed by the stimulation protocol described in the next subsection. The 561 nm line was used for ERK-KTR-mRuby2 and ERK-KTR-mCherry imaging (80% laser power; 150 ms), and 640 nm line for H2B-iRFP (80% laser power; 200 ms). OptoFGFR was stimulated and imaged with 488 nm (100% laser power, i.e., 1.15 W/cm^2^; 100 ms) on indicated time points.

#### Stimulation sequence

In all experiments, cells were stimulated with an identical sequence of light pulses (Table 1). The sequence was constructed as follows. First, based on preliminary data and earlier results (Nałęcz-Jawecki *et al*, 2023), we decided that the intervals between subsequent light pulses should approximately follow a Gamma distribution with shape *α* = 4 and scale *θ* = 5min. This choice was expected to be near-optimal in terms of bitrate and broad enough to probe various interval lengths. The actual number of intervals of a particular length in the sequence (Fig 1I) was chosen to best reflect this distribution given the time budget (∼17 h). Minor adjustments were made to ensure that each short interval length occurs at least twice. The intervals were ordered in such a way that the occurrences of a particular interval length were preceded by intervals maximally representative of the assumed distribution. For example, there were three intervals of length 13 min in the sequence; they were preceded by intervals of lengths 7, 17, and 28 min, which were close to, respectively, the 1st, 2nd, and 3rd quartile of the assumed distribution. This was important for a fair comparison between interval lengths (Fig 4) because a pulse following a short interval evokes a stronger response if the interval before it was also short; consequently, if all intervals of a particular length were preceded by short intervals, detectability of this interval length would be overestimated.

**Table 1.**
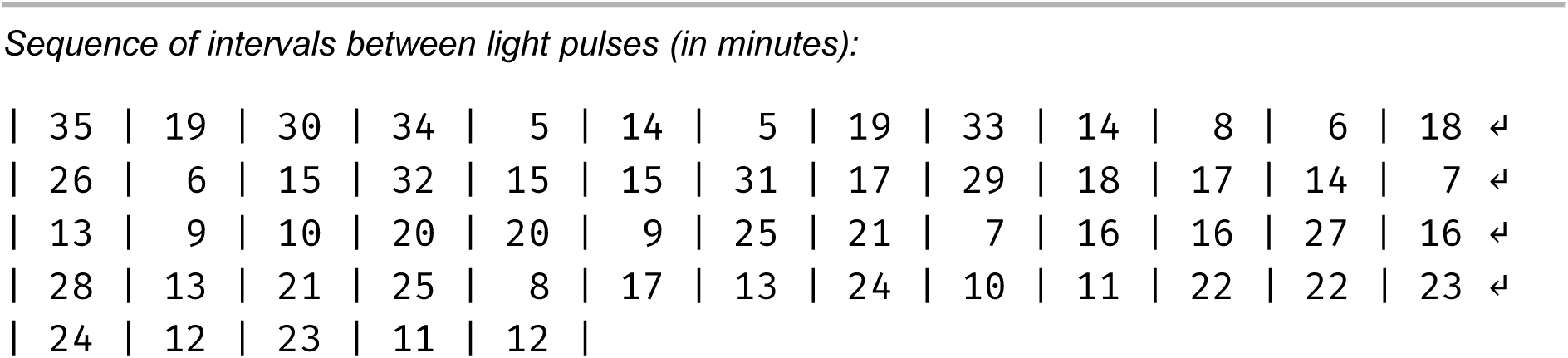
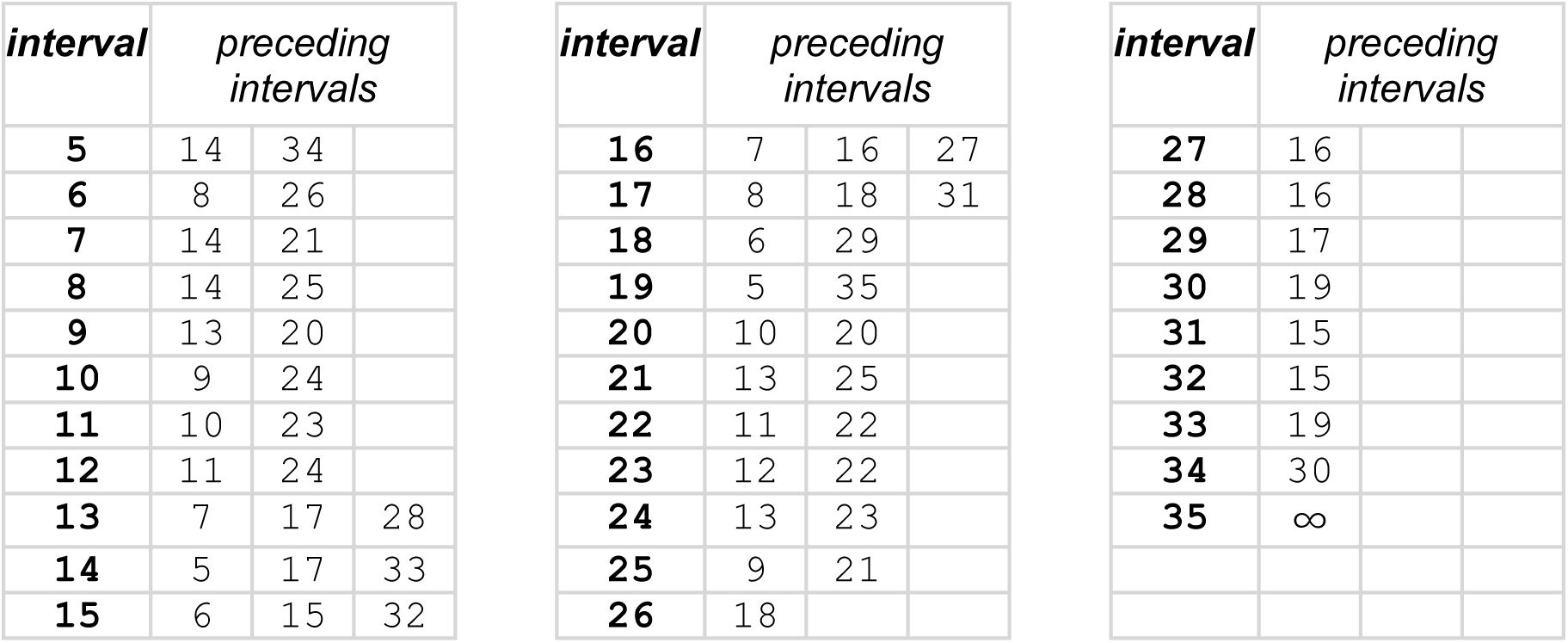
Sequence of light pulses used in experiments.

#### Image processing

Cell nuclei were detected based on H2B-iRFP fluorescence and tracked using our in-house software ShuttleTracker. Tracks shorter than 3 hours were removed from further analysis.

We estimated the background intensity in each channel as the mode of the pixel intensity (a common value was chosen across all frames and replicates with equal microscope settings) and subtracted it from the image. For each cell in a frame, we measured the average nuclear intensity in the ERK-KTR channel and normalized it with the average intensity over the whole frame. These two steps were necessary to eliminate global changes in mRuby2 fluorescence, arising in response to the 488 nm light used for optoFGFR stimulation, and to compensate for illumination differences across frames and replicates. The time course of the negative logarithm of the obtained value is referred to as a normalized *ERK-KTR trajectory*. We used this value instead of the typically used cytoplasmic to nuclear ratio to avoid noise associated with inaccurate cytoplasm detection. For clearer graphical presentation in Fig 2A, the single-cell ERK-KTR trajectories were normalized by subtracting the average value in each track.

We sorted the tracked nuclei in each replicate based on the mean optoFGFR fluorescence intensity in the time points in which optoFGFR was stimulated, and discarded tracks outside the range [*μ*, *μ* + 3*σ*] (BEAS-2B) or [*μ* − 0.5*σ*, *μ* + 2*σ*] (STE-1), where *μ* and *σ* denote the track-length-weighted average and standard deviation across all tracks in the particular replicate (see Fig S1).

For each track and light pulse, we computed the *response amplitude* as the log-ratio of ERK-KTR trajectory 7 min after pulse and at pulse. To account for the influence of the previous pulse, we normalized it by subtracting the average log-ratio of ERK-KTR trajectory (*L* + 7 min) and *L* after the pulse, where *L* is the interval between this and the previous pulse. The average was taken across all tracks and all pulses followed by at least (*L* + 7 min) interval (Fig S2).

We also computed the cell’s *responsiveness* as the standard deviation across all time points (not just time points with pulses) of the log fold change of translocation during 7 minutes. This measure was independent of pulse timing and, as such, could be used as a hint in automated pulse detection, as described below.

### Bitrate estimation

#### Derivation of the bitrate lower bound

We consider random sequences of light pulses that may occur in whole minutes. These sequences are encoded as *X* = *X*_1_, …, *X_N_* where *X_k_* = 1 if there was a pulse sent at the *k*-th minute and *X_k_* = 0 otherwise. We note that *X_k_* need not be independent. For each cell, we denote the sequence of responses (ERK-KTR trajectories) as *Y* = *Y*_1_, …, *Y_N_*. The mutual information between these two sequences can be computed as:

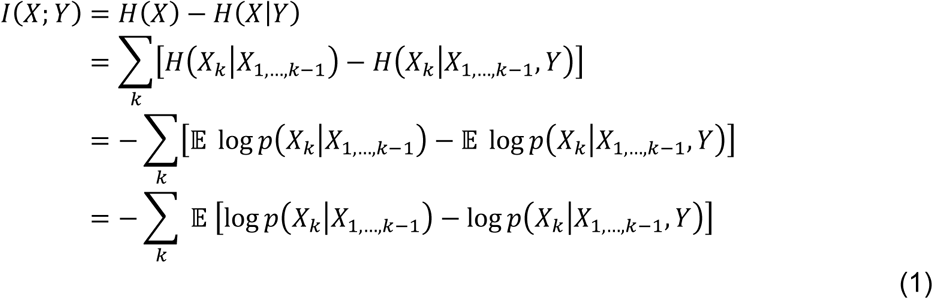

For independent *X_k_*, we would have *H*(*X_k_* | *X*_1…*k*−1_) = *H*(*X_k_*). In this case, intervals between pulses would follow a geometric distribution. However, we consider arbitrary interval distributions, and thus *H*(*X_k_* | *X*_1…*k*−1_) ≤ *H*(*X_k_*).

For a sufficiently large number of time points, *N*, the bitrate (or transmitted information per time point), b(*X*; *Y*), can be estimated as:

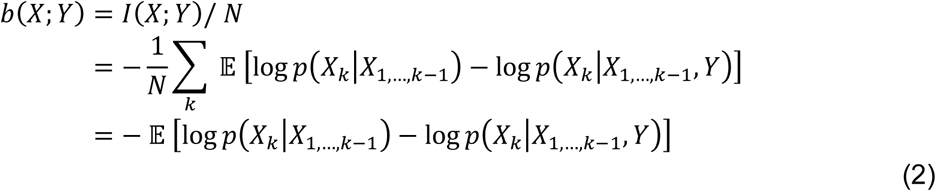

where, in the last equation, the expected value is also calculated over all time points *k* = 1, …, *N*. Thus, b(*X*; *Y*) is equal to the expected difference between the surprisal of *X_k_* conditioned on all previous pulses, −log p(*X_k_* | *X*_1…*k*−1_), and the surprisal of *X_k_* conditioned on all previous pulses and additionally the full response Y, −log p(*X_k_* | *X*_1…*k*−1_, *Y*). This can be thought of as the expected reduction in surprisal from learning *Y*.

To estimate b(*X*; *Y*), we first note that the probability p(*X_k_* | *X*_1…*k*−1_) depends only on the assumed encoding protocol. We restrict our analysis to protocols for which the probability of a pulse at time point *k* depends only on the time that elapsed since the previous pulse, *Last*_k_ (*Last*_k_ = *l* if and only if *X_k_*_−_*_l_* = 1 and *X_k_*_’_ = 0 for all *k*’ in {*k* − *l* + 1, …, *k* − 1}). Thus:

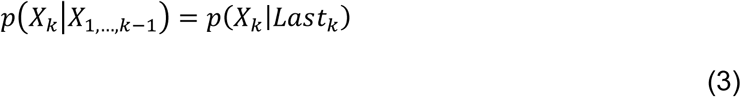

The posterior probability p(*X_k_* | *X*_1…*k*−1_, *Y*) cannot be directly computed. Instead, we estimate it by training a neural network p_*θ*_ (with weights *θ*) to predict *X_k_*. In principle, the neural network could use the complete stimulation history *X*_1…*k*−1_ and the full response *Y* to predict *X_k_*. However, we simplify this by reducing *X*_1…*k*−1_ to *Last_k_*, and *Y* to a slice *Y_k_*_…*k*+*r*_ (of length *r* + 1 = 7, starting at the time point *k*) and a scalar measure of cell responsiveness *S* = Std(*Y_k_* − *Y_k_*_+7_). This reduction is necessary as the network is trained and tested on the same sequence of pulses, and thus the stimulation history *X*_1…*k*−1_, as well as the full response trajectory *Y*, could be used to infer the timepoint in the experiment (*k*), and consequently reveal X_k_. By providing the network with *Last*_k_ instead of *X*_1…*k*−1,_ and the slice *Y_k_*_…*k*+*r*_ instead of *Y*, we prevent this leak. In summary, we approximate the posterior probability as:

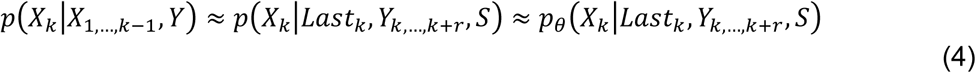

This approximation results in an upper bound on the conditional entropy:

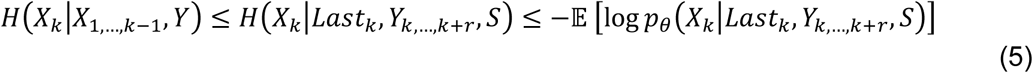

where the expectation is still computed over the true distribution of X, Y, and time points *k*. The first inequality follows from the data processing inequality, and the second reflects the fact that cross-entropy between p and p_*θ*_ is an upper bound on entropy of p. Thus, by substituting (5) back into formula (2), we get a lower bound on the bitrate:

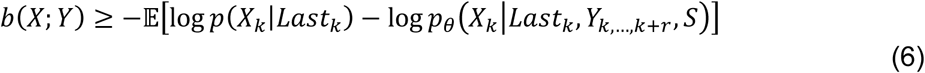

The bound is tight assuming the network correctly predicts the true posterior probability (both approximations in equation (4) are equalities). We estimate the bitrate by averaging over all cells *j* and timepoints *k* available in our dataset:

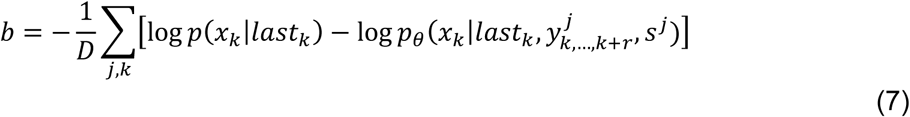

where *D* is the number of data points (*j*, *k*), and small letter symbols *x_k_* and *last*_k_ denote values from distributions *X_k_* and *Last*_k_.

#### Single-cell bitrate / Fraction of transmitting cells

We define the bitrate of a given cell track *j* as:

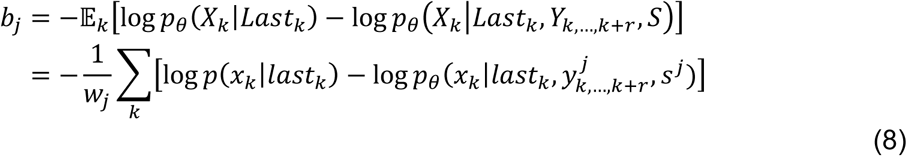

where *w_j_* is the number of timepoints available for cell track *j*. Note that since we use a model trained on all tracks, if a cell responds aberrantly, it can cause the model to make worse predictions than the prior, resulting in a negative bitrate of that track. The overall bitrate can be expressed as an average over the individual cell track bitrates (weighted by cell track length):

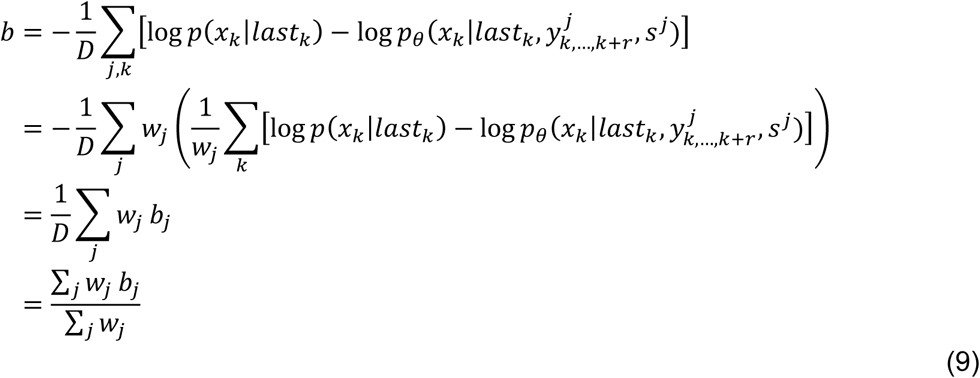

and since *w_j_* is the number of timepoints available for cell *j*, *D* = Σ*_j_ w_j_*.

The fraction of transmitting cells is determined by sorting the single-cell bitrates in ascending order and taking the longest prefix that averages (weighted by cell track length) to a negative value. Cells in the prefix are called non-transmitting, while the rest constitute the transmitting subpopulation.

### Training procedure

#### Data sampling

The network is trained to minimize the cross-entropy loss function using stochastic gradient descent:

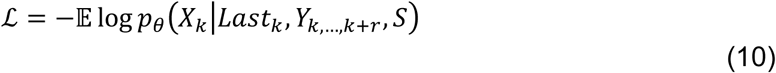

At each step, the expected value is computed by averaging over a mini-batch of 10^4^ data points (x_k_, *last*_k_, 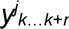, *s^j^*) randomly drawn from the dataset. In general, the network predictions *p*_*θ*_ depend on the assumed input protocol (interval distribution). Although throughout the paper we use a network trained according to the experimentally tested interval distribution, we designed the sampling procedure to allow for training and evaluation using an arbitrary protocol. We group all data points by *interval*_k_ (length of the interval containing the data point) and *last*_k_ (time elapsed since the previous pulse). We first sample pairs (*interval*_k_, *last*_k_) according to their distribution in the assumed protocol, then randomly retrieve data points from the corresponding groups in the experimental dataset to form mini-batches.

Procedure:

1. Group dataset by (*interval*_k_, *last*_k_).
2. Sample pairs of (*interval*_k_, *last*_k_) from the joint distribution of the assumed protocol.
3. Select random samples from the dataset for each (*interval*_k_, *last*_k_) pair.

#### Multiple experiments

Throughout the paper, we use a single network trained on all experiments. We allow the network to adapt to different experimental settings by supplying additional inputs:

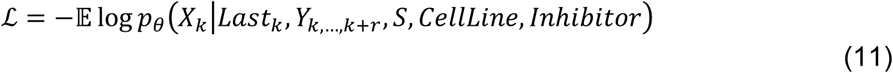

where *CellLine* is 0 for STE1 and 1 for BEAS-2B, and *Inhibitor* is a triplet of real numbers providing concentrations of 3 inhibitors {*ALKi, MEKi, CALCi*}. We then performed leave-one-out cross-validation on experimental replicates, see Fig S5. Since the change in bitrate estimate after exclusion of any single replicate is negligible, we concluded that the network does not overfit to individual replicates and we may use a single network trained on all available data.

#### Network architecture

The network inputs are log(*last*_k_), 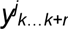, s*^j^ cellLine*, and *inhibitor* (3-dim). The trajectory slice 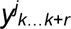 is replaced with discrete differences 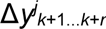, where 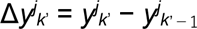 to abstract from the base level of ERK-KTR expression in individual cells. We chose to use the slice of *r* = 6 discrete differences, as its further extension provides no substantial increase in bitrate. All inputs are concatenated, resulting in a feature vector of length (*r* + 6). The features are standardized across the dataset to have zero mean and a standard deviation of *σ* = 1, and passed to a three-layer perceptron (MLP) with hidden layer sizes of (40, 20) and leaky ReLU activations. The MLP computes a logit (Bayesian) update *u*_Bayes_ to the prior probability *p*(*X_k_* = 1 | *Last*_k_):

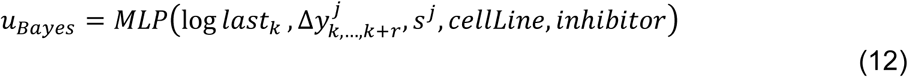

This update is then used to compute the posterior (logit) probability of X_k_ = 1:

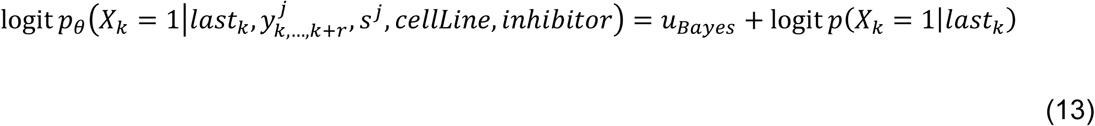

This approach allows the network to generalize more effectively across protocols with different priors *p*(*X_k_* = 1 | *Last*_k_), simplifying protocol optimization.

Fig S6 illustrates the mean updates u_Bayes_ generated by the MLP, grouped by (*interval*_k_, *last*_k_) for STE-1 with ALKi. A positive value indicates that the network consistently predicts a higher pulse probability than the prior—this is the case along the diagonal (*interval*_k_ = *last*_k_), corresponding to timepoints with actual pulses (*X_k_* = 1). A value below zero indicates that the network predicts a lower pulse probability than the prior. A value close to zero implies that the network cannot improve upon the prior probability. As a result, no information is gained. This behavior is observed for *last*_k_ ≤ 10, where cells have not yet recovered from a previous pulse and cannot respond to a new pulse (see also Fig 4A). Consequently, their ERK-KTR trajectory carries no information about a potential new pulse. For *last*_k_ > 10, off-diagonal terms (where *X_k_* = 0) are generally negative. For such *last*_k_, a pulse would evoke a noticeable ERK-KTR trajectory change, and thus the absence of such change leads the network to infer that no pulse has occurred and predict a probability lower than the prior. Importantly, these negative predictions also contribute to the total information transmitted, and thus the bitrate. Finally, we note that entries immediately next to the diagonal are also close to zero. This suggests that the network sees a response to the pulse, but is not perfectly sure of the exact timing of the pulse.

### Protocol optimization

To optimize the protocol, we want to directly *maximize* the bitrate estimate:

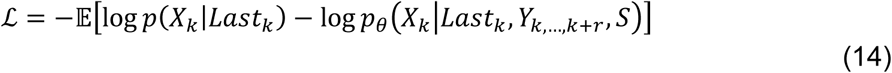

This introduces several challenges. First, during training, the protocol’s interval probabilities are optimized via gradient descent. However, the expected value is computed by sampling according to the protocol, and thus we need to compute a derivative of the form *∇*_*ϕ*_ 𝔼*_z_*_∼*Z*(*ϕ*)_ f(*z*), for some distribution *Z* parametrized by *ϕ*. This is done by rewriting the expected value using a frozen distribution with importance weights:

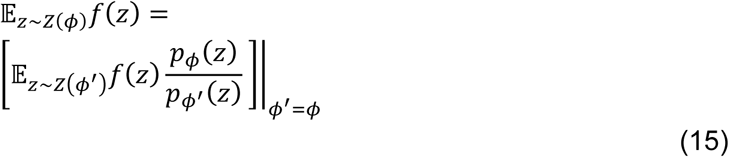

Since the sampled distribution no longer depends on *ϕ*, the gradient *∇*_*ϕ*_ may be calculated as follows:

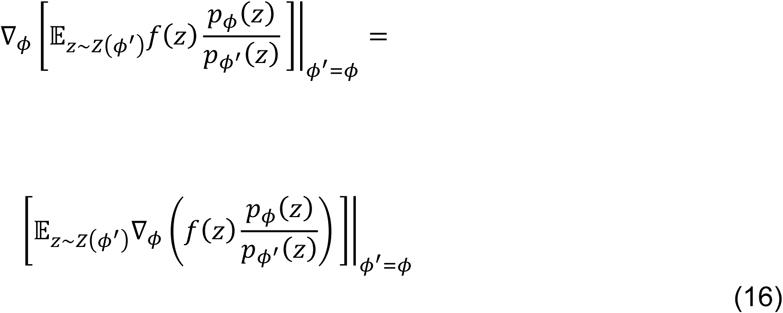

Although the importance weights *p*_*ϕ*_(*z*) / *p*_*ϕ*’_(*z*) are equal to 1, their derivatives with respect to *ϕ* in general differ from zero and allow us to compute the gradient. In our case, we sample from a frozen protocol and update the protocol after calculating each gradient step.

Secondly, the optimal protocol should assign non-zero probabilities to intervals longer than 35 min, the longest interval used in our experiments. Since we observe that responses for intervals longer than 30 min are similar (and strongests), we handle longer intervals by imputing them using the available 35-minute interval from our experiments. Specifically, when a trajectory slice for a pair (*interval*_k_, *last*_k_) with *interval*_k_ > 35 min is requested, we return a slice for an imputed pair (*interval*_k_*, *last*_k_*) instead, where *interval*_k_* = 35 min and *last*_k_* is determined as follows:

- If 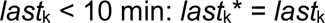
- If 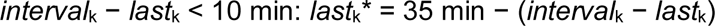
- Otherwise:

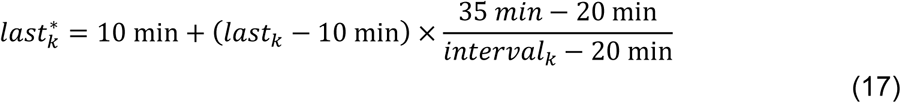

This way, we keep the time since the last pulse and the time to the next pulse fixed if this time is shorter than 10 min. A graphical illustration of imputation for *interval_k_* = 60 min is shown in Fig S7. Note that while *y_k_*_…*k*+*r*_ is imputed, the network is provided with true *last*_k_.

Furthermore, selection of the transmitting subpopulation technically depends on single-cell bitrates, and thus also on the protocol. We perform protocol optimization on the transmitting subpopulation determined using the experimental protocol.

Finally, the optimized protocol can exhibit large jumps in probabilities of specific intervals due to experimental noise and the order in which intervals were tested in the experiment. To address this, we introduce a regularization term in the loss function to promote smoother protocols:

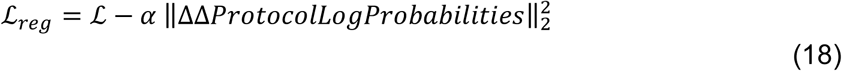

where ΔΔ*ProtocolLogProbabilities* represents the second-order differences of the interval log-probabilities, and 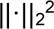 is the mean of squares. We found that setting *α* = 0.003 yields noticeably smoother protocols with minimal impact on the bitrate (see Fig 4C and Fig S8).

## Author contributions

Conceptualization: P.N-J., L.R., F.G., M.K., L.J.B., T.L.

Methodology: P.N-J., L.R., F.G., M.K., L.J.B., T.L.

Software: P.N-J., F.G., M.K.

Validation: P.N-J., L.R., M.K.

Formal analysis: P.N-J., F.G.

Investigation: P.N-J., L.R., F.G.

Resources: L.R., S.L., M.K., L.J.B.

Data curation: P.N-J., F.G., M.K.

Writing – original draft: P.N-J., F.G., L.J.B., T.L.

Writing – review & editing: P.N.-J., L.R, F.G., S.L., M.K., L.J.B., T.L,

Visualization: P.N-J., L.R., F.G., L.J.B

Supervision: L.J.B., T.L,

Project administration: P.N-J., L.J.B., T.L.

Funding acquisition: L.J.B., T.L.

## Data Availability

Data used in the paper (time-lapse microscopy quantifications) are deposited on Zenodo (https://doi.org/10.5281/zenodo.15282876). Raw images can be obtained from the authors. Source code is available on Github (https://github.com/pawelnalecz/pathway-bitrate).

## Appendix

**Fig S1.**
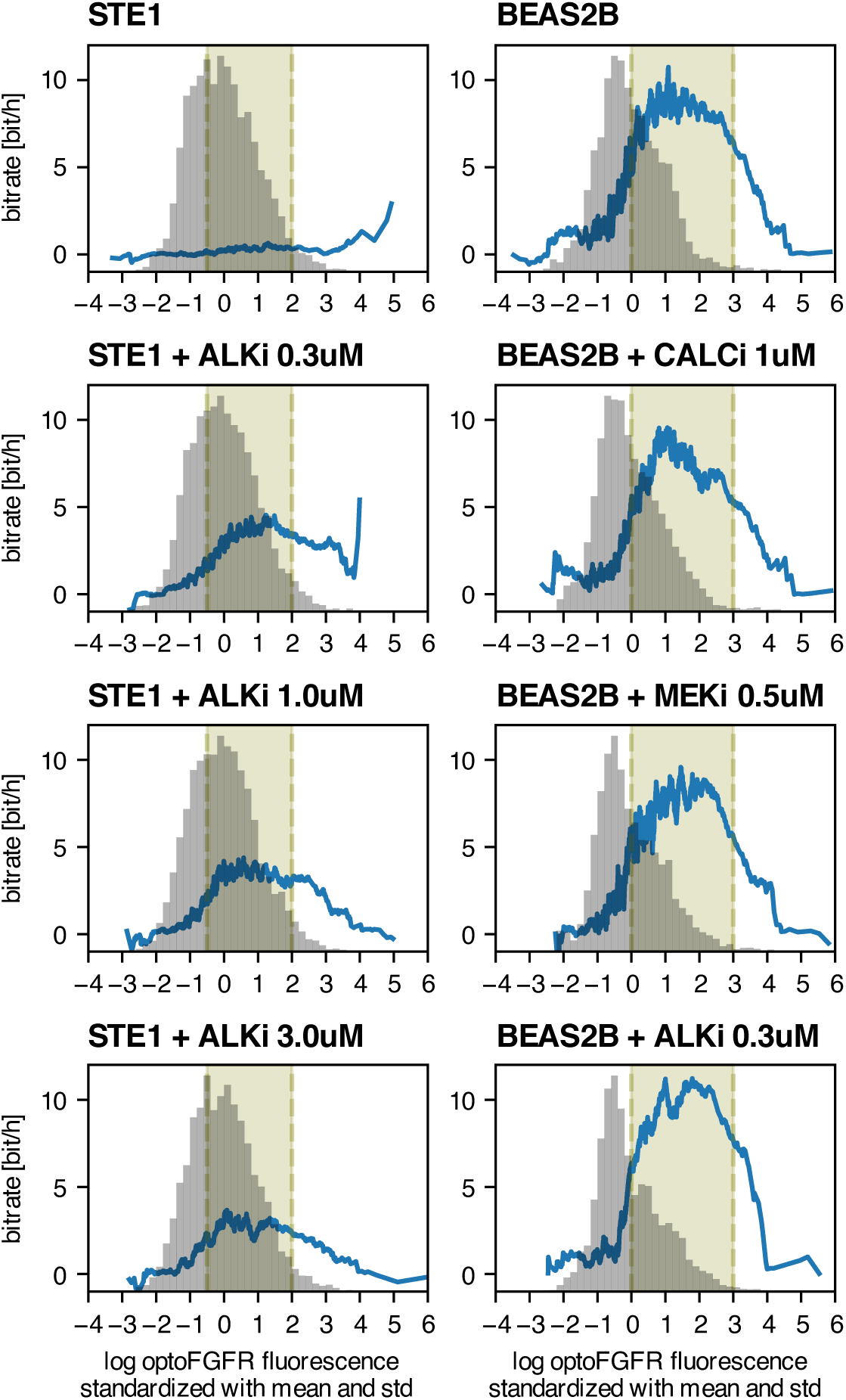
Cell preselection based on optoFGFR receptor level. Average bitrate in individual cells as a function of mean optoFGFR fluorescence (rolling mean over single-cell trajectories; trajectories in each rolling window have a combined duration equal to 100 times the experiment’s duration), superimposed over the histograms of mean optoFGFR fluorescence in individual cells. OptoFGFR levels were log-transformed and standardized within each replicate separately, then pooled by cell line and condition. Cells in the highlighted range were used in further analysis.

**Fig S2.**
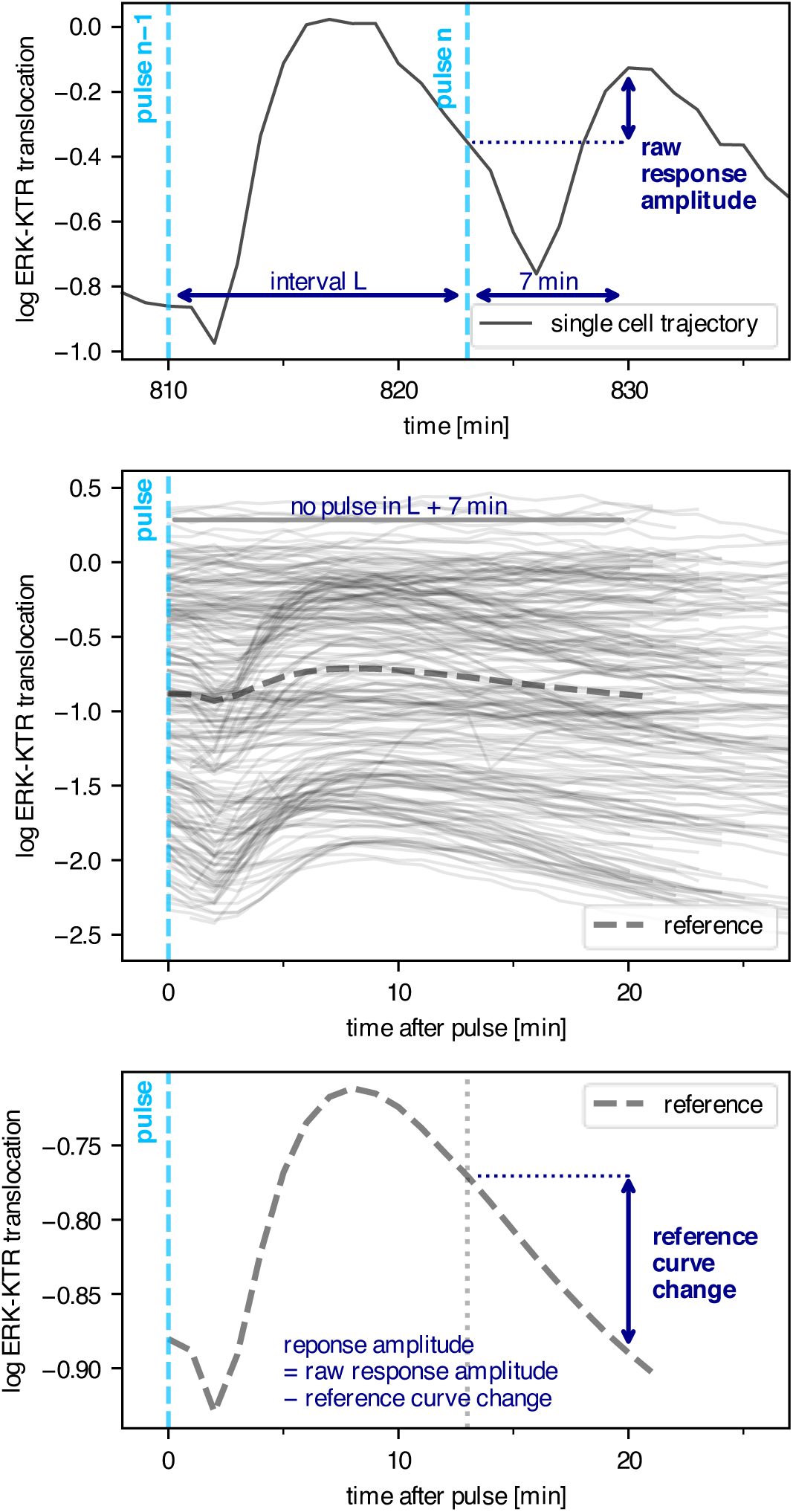
Response amplitude. (A) To compute the response amplitude to light pulse *n*, which occurred time *L* after pulse (*n* − 1), we first calculated the raw response amplitude as the difference between the ERK-KTR trajectory at pulse and 7 min after. (B) We constructed a reference trajectory by averaging the ERK-KTR trajectories after all pulses not followed by another pulse within the next *L* + 7 min. (C) We computed the change in the reference trajectory between time *L* and (*L* + 7 min) after the pulse. The response amplitude was calculated by subtracting the obtained value from the raw response amplitude.

**Fig S3.**
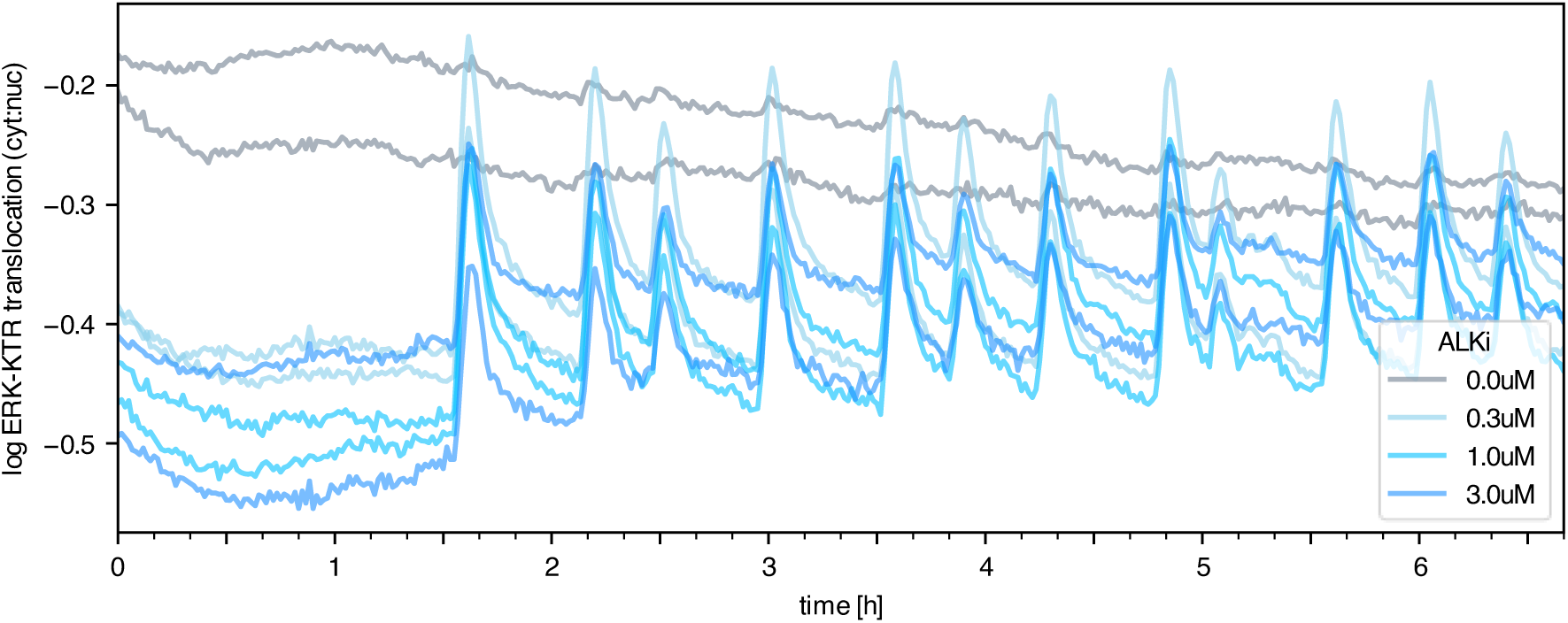
Comparison of ERK-KTR translocation in STE-1 cells with and without ALKi. Each line represents the ERK-KTR trajectory averaged over all cells in a single technical replicate. Note that in this figure (and only here), the ERK-KTR trajectory is computed as the log-ratio of cytoplasmic and nuclear ERK-KTR fluorescence, which allows for comparison between conditions and replicates.

**Fig S4.**
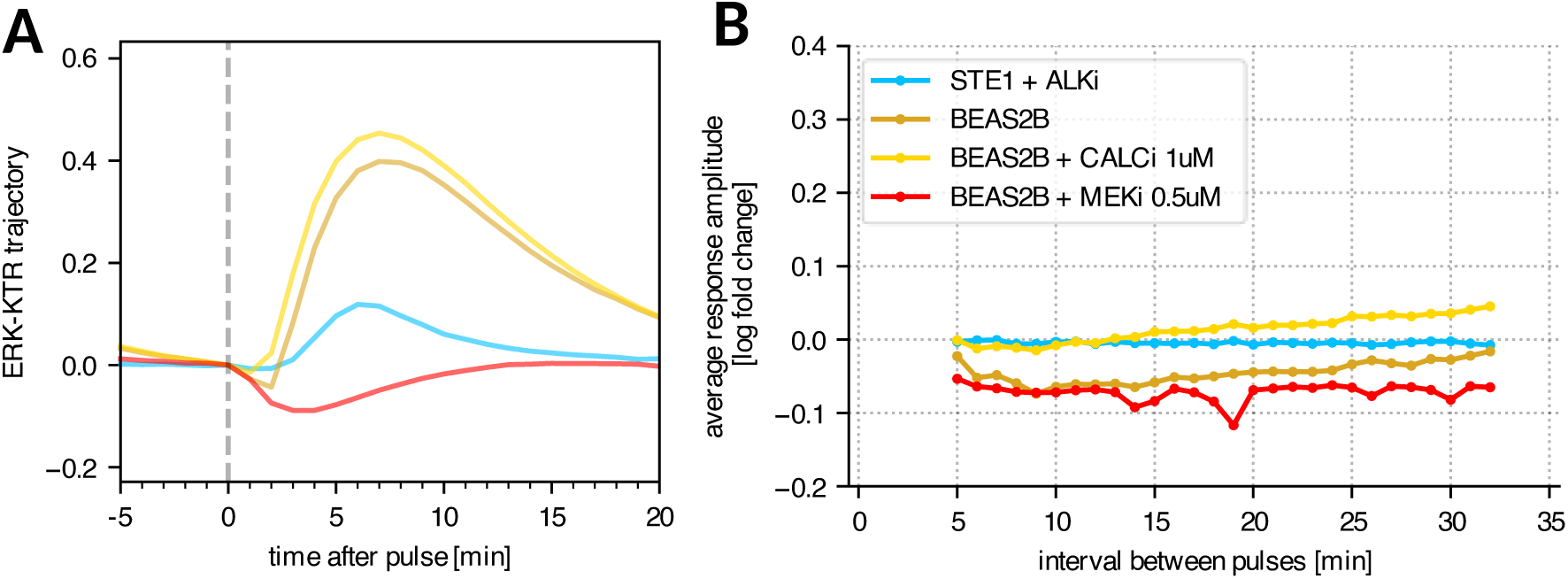
Dip amplitude. (A) Population-average ERK-KTR response to a sample stimulation pulse. Color lines correspond to cell lines and conditions, as shown in panel B. The ERK-KTR dip is visible only in BEAS-2B cells with no inhibitor and with MEK inhibitor. (B) Average response amplitude 2 min after light pulse as a function of time since the previous light pulse.

**Fig S5.**
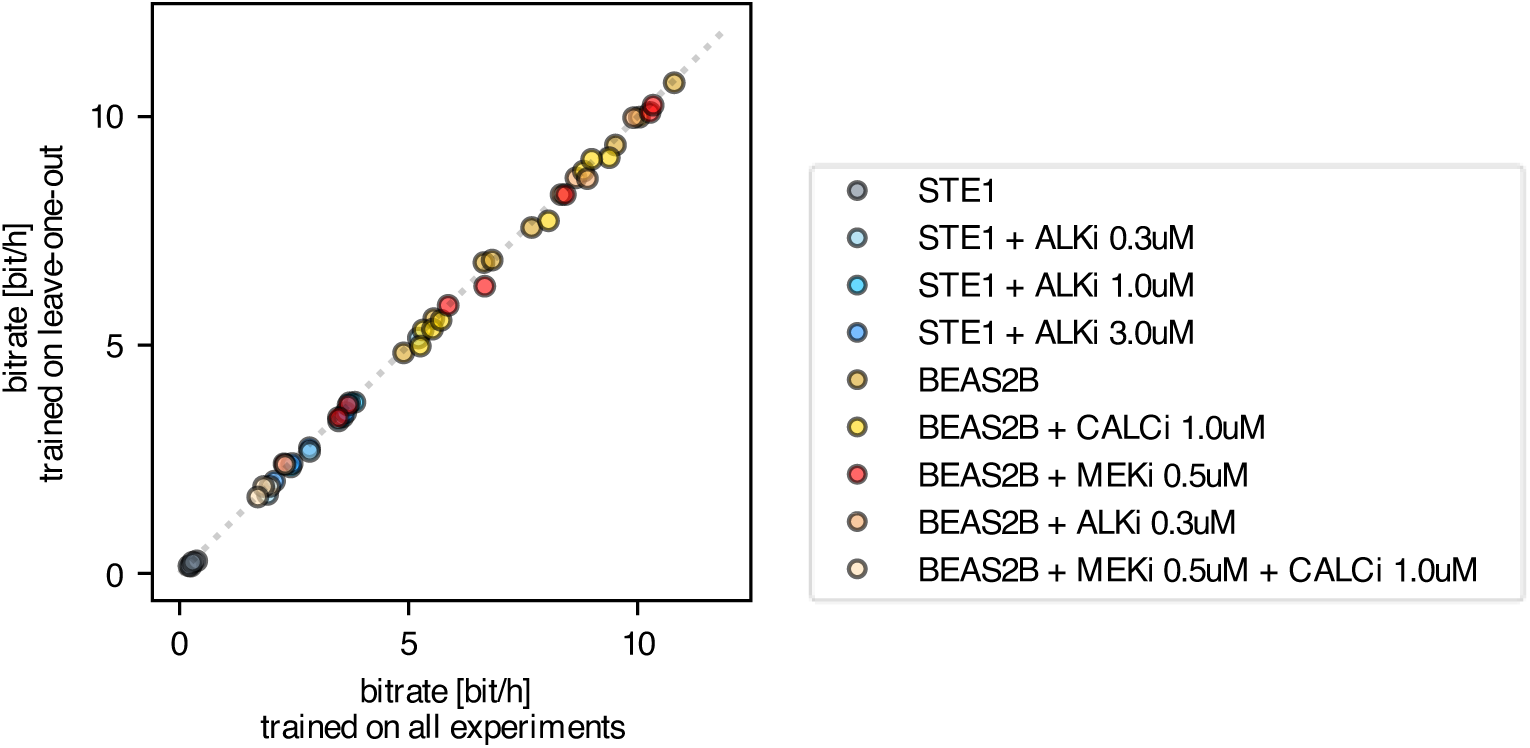
Leave-one-out network evaluation. Average bitrate in particular replicates, computed based on predictions by MLP trained on all conditions and replicates (as used throughout the paper; x-axis), and on all conditions and replicates except for the currently evaluated replicate (y-axis). Dotted line denotes the identity line.

**Fig S6.**
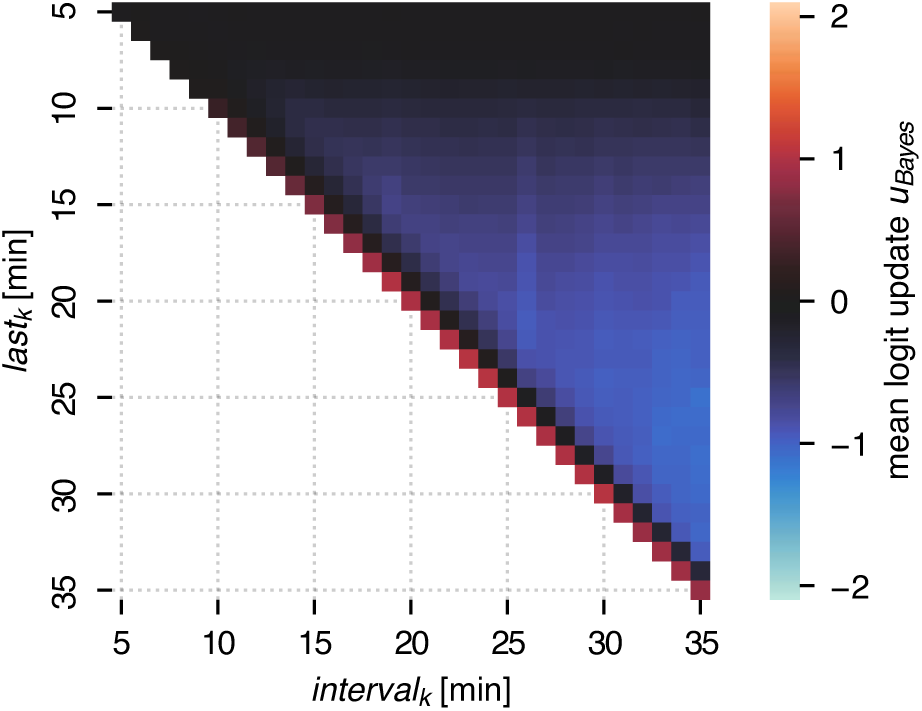
Mean logit Bayesian update *u*_Bayes_ as a function of *last*_k_ and *interval*_k_. evaluated on STE1 cells with ALKi. The diagonal (*last*_k_ = *interval*_k_) corresponds to timepoints in which a pulse occurred; the *u*_Bayes_ values on the diagonal are presented in Fig 4B.

**Fig S7.**
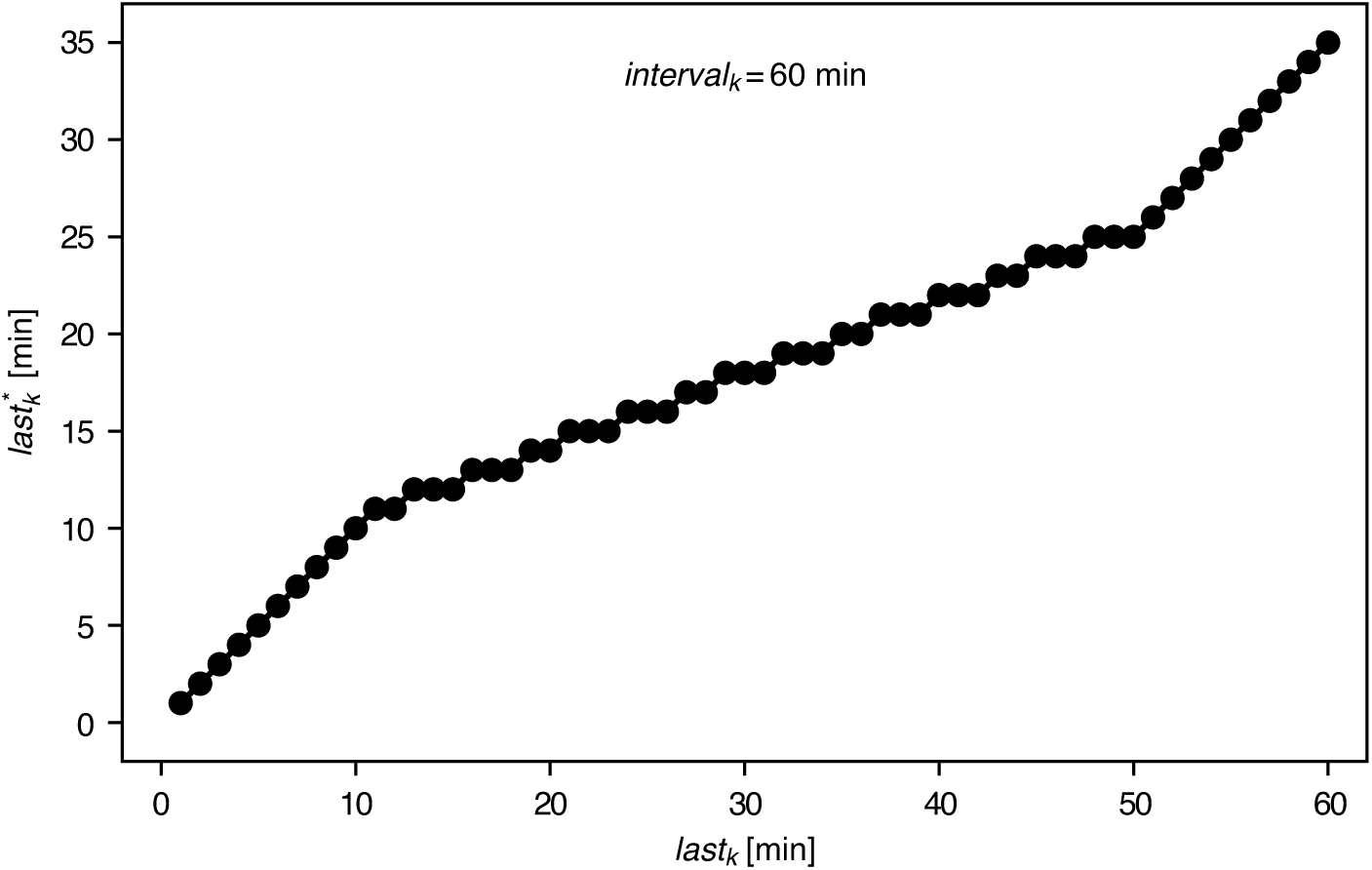
Imputation of responses after intervals not present in the dataset. Since we do not have experimental data for responses (*interval*_k_, *last*_k_) for *interval*_k_ > 35 min, we instead sample from responses (*interval*_k_* = 35 min, *last*_k_*). The plot illustrates the mapping of *last*_k_ to *last*_k_* in the case of *interval*_k_ = 60 min.

**Fig S8.**
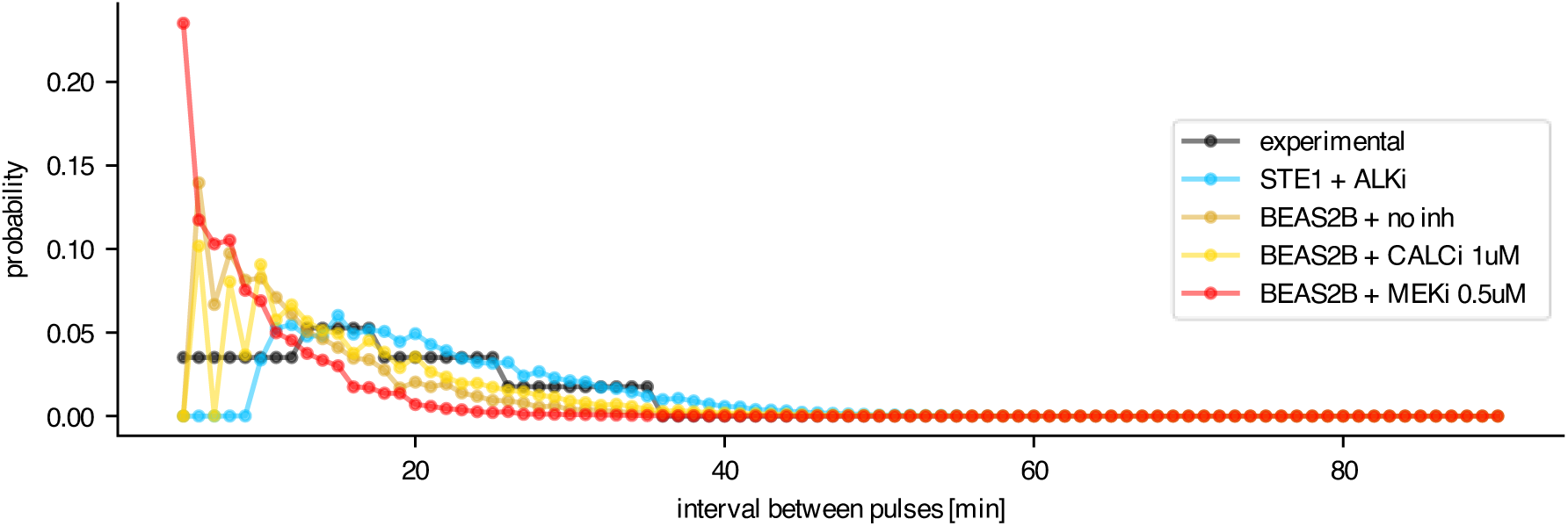
Input interval distribution optimized without regularization. Setup as in Fig 4C.

## References

Aoki K, Kondo Y, Naoki H, Hiratsuka T, Itoh RE & Matsuda M (2017) Propagating Wave of ERK Activation Orients Collective Cell Migration. Dev Cell 43: 305–317.e5

Bartkova J, Hořejší Z, Koed K, Krämer A, Tort F, Zieger K, Guldberg P, Sehested M, Nesland JM, Lukas C, et al (2005) DNA damage response as a candidate anti- cancer barrier in early human tumorigenesis. Nature 434: 864–870

Berridge MJ (1995) Calcium signalling and cell proliferation. BioEssays 17: 491–500

Bugaj LJ, Sabnis AJ, Mitchell A, Garbarino JE, Toettcher JE, Bivona TG & Lim WA (2018) Cancer mutations and targeted drugs can disrupt dynamic signal encoding by the Ras-Erk pathway. Science 361: eaao3048

Burotto M, Chiou VL, Lee J & Kohn EC (2014) The MAPK pathway across different malignancies: A new perspective. Cancer 120: 3446–3456

Cheong R, Rhee A, Wang CJ, Nemenman I & Levchenko A (2011) Information transduction capacity of noisy biochemical signaling networks. Science 334: 354–358

Corson LB, Yamanaka Y, Lai K-MV & Rossant J (2003) Spatial and temporal patterns of ERK signaling during mouse embryogenesis. Development 130: 4527–4537

DerMardirossian C (2024) The Role of Calcium in Actin-Dependent Cell Migration and Invasion in Cancer. In Physiology, Heinbockel T (ed) IntechOpen

Dessauges C, Mikelson J, Dobrzyński M, Jacques M, Frismantiene A, Gagliardi PA, Khammash M & Pertz O (2022) Optogenetic actuator – ERK biosensor circuits identify MAPK network nodes that shape ERK dynamics. Mol Syst Biol 18: e10670

Gerosa L, Chidley C, Fröhlich F, Sanchez G, Lim SK, Muhlich J, Chen J-Y, Vallabhaneni S, Baker GJ, Schapiro D, et al (2020) Receptor-Driven ERK Pulses Reconfigure MAPK Signaling and Enable Persistence of Drug-Adapted BRAF-Mutant Melanoma Cells. Cell Systems 11: 478–494.e9

Gilkey JC, Jaffe LF, Ridgway EB & Reynolds GT (1978) A free calcium wave traverses the activating egg of the medaka, Oryzias latipes. J Cell Biol 76: 448–466

Gonzalez-Martinez D, Roth L, Mumford TR, Guan J, Le A, Doebele RC, Huang B, Tulpule A, Niewiadomska-Bugaj M, Bivona TG, et al (2024) Oncogenic EML4-ALK assemblies suppress growth factor perception and modulate drug tolerance. Nat Commun 15: 9473

Grabowski F, Czyż P, Kochańczyk M & Lipniacki T (2019) Limits to the rate of information transmission through the MAPK pathway. J R Soc Interface 16: 20180792

Hino N, Rossetti L, Marín-Llauradó A, Aoki K, Trepat X, Matsuda M & Hirashima T (2020) ERK-Mediated Mechanochemical Waves Direct Collective Cell Polarization. Dev Cell 53: 646–660.e8

Katso R, Okkenhaug K, Ahmadi K, White S, Timms J & Waterfield MD (2001) Cellular Function of Phosphoinositide 3-Kinases: Implications for Development, Immunity, Homeostasis, and Cancer. Annu Rev Cell Dev Biol 17: 615–675

Kim N, Kim JM, Lee M, Kim CY, Chang K-Y & Heo WD (2014) Spatiotemporal control of fibroblast growth factor receptor signals by blue light. Chem Biol 21: 903–912

Kramar M, Hahn L, Walczak AM, Mora T & Coppey M (2024) Single cells can resolve graded stimuli. doi:10.1101/2024.10.29.620645 [PREPRINT]

Kramer BA, Sarabia Del Castillo J & Pelkmans L (2022) Multimodal perception links cellular state to decision-making in single cells. Science 377: 642–648

Lee KW, Moreau M, Néant I, Bibonne A & Leclerc C (2009) FGF-activated calcium channels control neural gene expression in Xenopus. Biochimica et Biophysica Acta (BBA) - Molecular Cell Research 1793: 1033–1040

Lei Y, Lei Y, Shi X & Wang J (2022) EML4-ALK fusion gene in non-small cell lung cancer (Review). Oncol Lett 24: 277

Marei HE, Althani A, Afifi N, Hasan A, Caceci T, Pozzoli G, Morrione A, Giordano A & Cenciarelli C (2021) p53 signaling in cancer progression and therapy. Cancer Cell Int 21: 703

Mehrdad AE & Parvin M (2018) ATM in breast and brain tumors: a comprehensive review. Cancer Biology & Medicine 15: 210

Nałęcz-Jawecki P, Gagliardi PA, Kochańczyk M, Dessauges C, Pertz O & Lipniacki T (2023) The MAPK/ERK channel capacity exceeds 6 bit/hour. PLOS Computational Biology 19: e1011155

Nguyen HB, Estacion M & Gargus JJ (1997) Mutations Causing Achondroplasia and Thanatophoric Dysplasia Alter bFGF-Induced Calcium Signals in Human Diploid Fibroblasts. Human Molecular Genetics 6: 681–688

Nishida E & Gotoh Y (1993) The MAP kinase cascade is essential for diverse signal transduction pathways. Trends in Biochemical Sciences 18: 128–131

Selimkhanov J, Taylor B, Yao J, Pilko A, Albeck J, Hoffmann A, Tsimring L & Wollman R (2014) Accurate information transmission through dynamic biochemical signaling networks. Science 346: 1370–1373

Shannon CE (1948) A Mathematical Theory of Communication. Bell Syst Tech J 27: 379– 423

Soda M, Choi YL, Enomoto M, Takada S, Yamashita Y, Ishikawa S, Fujiwara S, Watanabe H, Kurashina K, Hatanaka H, et al (2007) Identification of the transforming EML4– ALK fusion gene in non-small-cell lung cancer. Nature 448: 561–566

Song Y, Bi Z, Liu Y, Qin F, Wei Y & Wei X (2023) Targeting RAS–RAF–MEK–ERK signaling pathway in human cancer: Current status in clinical trials. Genes & Diseases 10: 76– 88

Sugimoto T, Stewart S & Guan K-L (1997) The Calcium/Calmodulin-dependent Protein Phosphatase Calcineurin Is the Major Elk-1 Phosphatase. Journal of Biological Chemistry 272: 29415–29418

Tkačik G, Callan CG & Bialek W (2008) Information flow and optimization in transcriptional regulation. Proc Natl Acad Sci USA 105: 12265–12270

Topolewski P, Zakrzewska KE, Walczak J, Nienałtowski K, Müller-Newen G, Singh A & Komorowski M (2022) Phenotypic variability, not noise, accounts for most of the cell-to-cell heterogeneity in IFN-γ and oncostatin M signaling responses. Sci Signal 15: eabd9303

Tudelska K, Markiewicz J, Kochańczyk M, Czerkies M, Prus W, Korwek Z, Abdi A, Błoński S, Kaźmierczak B & Lipniacki T (2017) Information processing in the NF-κB pathway. Sci Rep 7: 15926

Uda S, Saito TH, Kudo T, Kokaji T, Tsuchiya T, Kubota H, Komori Y, Ozaki Y & Kuroda S (2013) Robustness and compensation of information transmission of signaling pathways. Science 341: 558–561

Whitaker M (2006) Calcium at Fertilization and in Early Development. Physiological Reviews 86: 25–88

Zhang W & Liu HT (2002) MAPK signal pathways in the regulation of cell proliferation in mammalian cells. Cell Res 12: 9–18

